# *C. elegans* Dicer stacks with the RIG-I-like receptor DRH-1 to cleave dsRNA

**DOI:** 10.64898/2026.08.16.745104

**Authors:** Claudia D. Consalvo, Parker J. Nichols, Elaina P. Boyle, P. Joseph Aruscavage, Peter S. Shen, Brenda L. Bass

## Abstract

In prior studies we showed that the nematode ancestor of Dicer’s helicase domain had minimal ATP hydrolysis, translocation and dsRNA binding activity, and in extant *C. elegans* the RIG-I like receptor (RLR) DRH-1, and the dsRNA binding protein RDE-4, were co-opted to provide these activities. Here we report cleavage-competent cryo-EM structures of the antiviral complex (AVC; DCR-1•DRH-1•RDE4), in the absence (3.2Å) and presence (3.0Å) of ATP. A key feature of both structures is stacking of DCR-1’s helicase with the helicases of two DRH-1 molecules, reminiscent of oligomerization of mammalian RLRs. dsRNA threads through all three helicases with a widened major groove at helicase-helicase interfaces and where the conserved Hel2 loop inserts into the major groove. The presence of nucleotide redistributed helicase– helicase and helicase–RNA contacts in the AVC, as for MDA5, albeit specific interactions and remodeling differed. MDA5 and RIG-I RLRs contain an unstructured linker between their CARDs and helicase domain, but the analogous linker in DRH-1 has a short-structured region positioned to interact with DCR-1. These findings provide a structural framework for understanding how DCR-1, DRH-1, and RDE-4 cooperate to cleave viral dsRNA and elucidate unique features required for antiviral defense in different animals.

## INTRODUCTION

Dicer is a multidomain RNase III enzyme with an essential role in generating small regulatory RNAs in eukaryotes^1^. Like all RNase III enzymes, Dicer’s cleavage activity is mediated by conserved RNase III domains. Simpler RNase III enzymes in bacteria contain a single RNase III domain and function as dimers. In contrast, Dicer has two RNase III domains (RNase IIIa and RNase IIIb) within a single polypeptide that form an intramolecular dimer to simultaneously cleave both strands of dsRNA^1–3^. Cleavage of dsRNA by Dicer generates small regulatory RNAs, such as microRNAs (miRNAs) and small interfering RNAs (siRNAs), that are typically 21–27 nucleotides (nts) long with 2nt 3′ overhangs (ovrs)^3–5^. In addition to its two RNase III domains, Dicer possesses other conserved regions, including an N-terminal helicase domain, a DUF283 domain, a PAZ domain that binds the 3′ovrs of dsRNA, and a C-terminal dsRNA-binding motif (dsRBM; Figure 1A)^6,7^.

**Figure 1.**
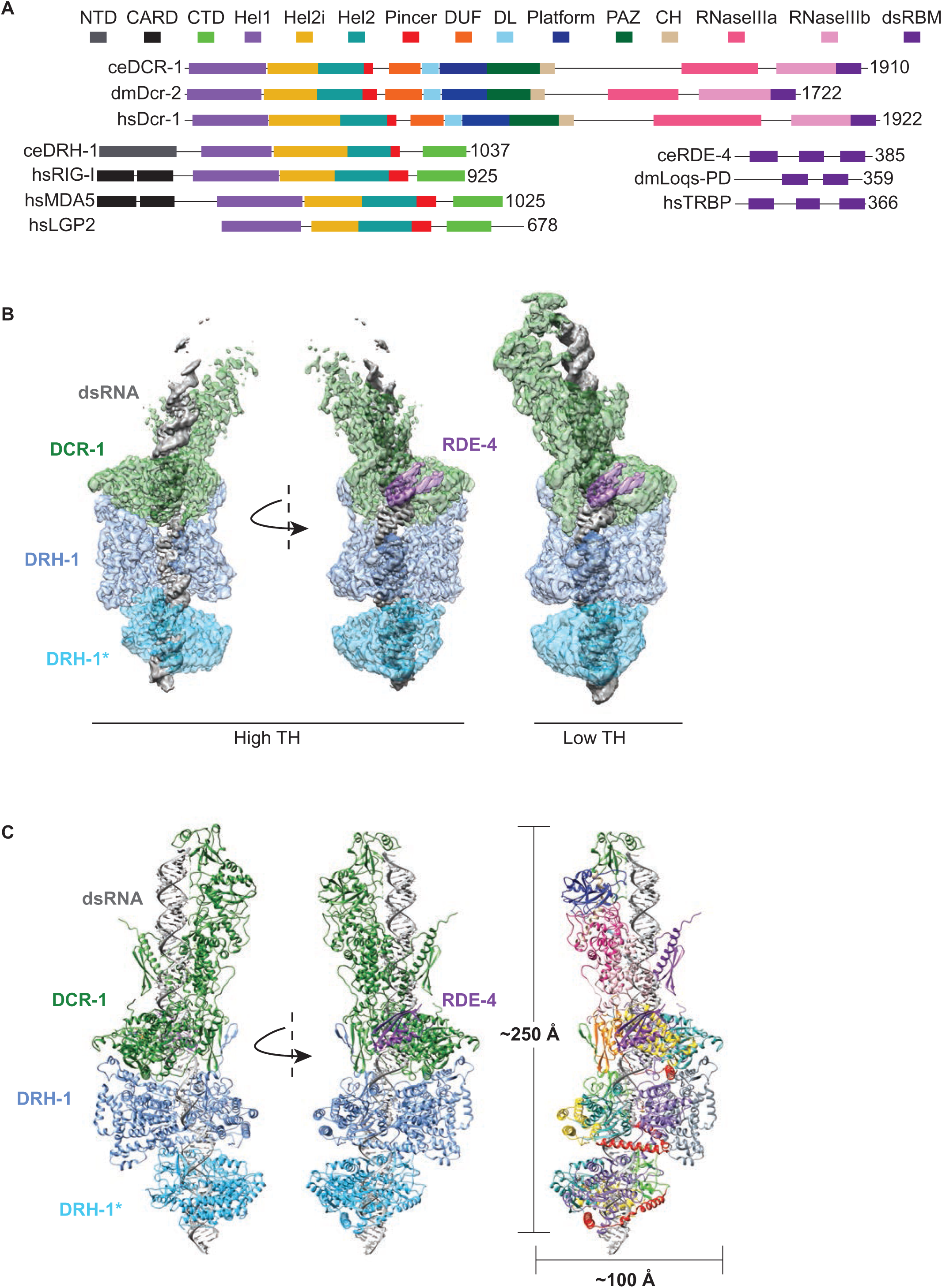
Overall structure of the AVC in a cleavage-competent state. A. Colored rectangles depict conserved domains and are adapted from *Consalvo et al*^20^. Numbers to right of open-reading frame indicate total amino acids in protein. Organism is indicated: ce, *C. elegans*; dm, *Drosophila melanogaster,* hs, *Homo sapiens*. Dicer domains DL (DUF-Linker) and Connecter Helix (CH) are indicated, but not discussed further. B. Cryo-EM reconstruction of the AVC at 3.0Å resolution. DCR-1 is green, DRH-1 is blue (different shades represent individual monomers), RDE-4 is purple, and dsRNA is gray_._ DRH-1* refers to only the helicase and CTD domains being resolved. Two thresholds (TH) are displayed for clarity. Protein densities were made transparent to better visualize dsRNA threaded through all three helicase domains. C. Model of the AVC complex. On left, each protein is colored as in (B). On right, domains of each protein are colored as depicted in (A).

Dicer’s multidomain architecture and its interaction with other dsRNA binding proteins (dsRBPs) allow it to recognize, bind, and accurately process dsRNA substrates^7,8^. The resulting small RNA products are then incorporated into the RNA-induced silencing complex (RISC), which guides the small RNAs to the complementary RNA targets to mediate their degradation or translational repression, a process known as RNA interference (RNAi)^9^. miRNAs primarily regulate endogenous gene expression, whereas siRNAs are guided to foreign genetic elements like viruses and transposons^9^. Through these pathways, Dicer plays a critical role in controlling gene expression, maintaining genome stability, and defending against viral infections^10^.

While Dicer performs a conserved core function of processing dsRNA into small regulatory RNAs, its specific roles and mechanisms vary among organisms^7,8^. Many invertebrates encode multiple Dicer enzymes with specialized roles. For example, *Drosophila melanogaster* has two Dicers—dmDcr1 and dmDcr2. dmDcr1 specializes in miRNA biogenesis for gene regulation, whereas dmDcr2 produces siRNA from exogenous long dsRNAs, including those associated with viral replication or transposons^11–14^. In contrast, vertebrates typically have a single Dicer enzyme devoted to miRNA processing. Antiviral defense is instead mediated by RIG-I-like receptors (RLRs)^15^. RLRs are a family of cytoplasmic pattern recognition receptors that detect viral dsRNA and activate the interferon response. Despite these divergent antiviral strategies, Dicer and RLRs share an evolutionarily conserved helicase domain that plays a critical role in RNA sensing^15,16^.

In *C. elegans*, the sole Dicer enzyme (DCR-1) functions across several RNAi pathways by forming distinct protein complexes with different types of dsRNA. For antiviral defense, DCR-1 functions with the dsRBP RDE-4 and the RLR ortholog DRH-1^17–19^. It is common to see Dicer enzymes interact with dsRBPs similar to RDE-4, where these proteins contain two or three dsRBMs and regulate or enhance Dicer cleavage activity^8^. In contrast, the interaction between DCR-1 and DRH-1 is the only known example of Dicer and RLRs working together to promote antiviral defense. We previously determined two low-resolution structures of the *C. elegans* antiviral complex (AVC; DCR-1•DRH-1•RDE-4) that suggested it is the N-terminal domain (NTD) of DRH-1 that interacts with DCR-1’s helicase domain. However, neither structure represented a cleavage-competent state. Both structures showed dsRNA bound to the helicase domain of DRH-1, but in one structure there was a second dsRNA bound to DCR-1 that was positioned away from the RNase III domains^20^.

Here we used cryo-EM to determine how the *C. elegans* AVC cleaves viral dsRNA. By optimizing sample conditions, we obtained 3.0Å and 3.2Å reconstructions of the AVC in the presence and absence of ATP, respectively. Both structures revealed a cleavage-competent conformation with dsRNA threaded through three stacked helicase domains: one from DCR-1 and two from the two additional stacked DRH-1 molecules. Because this architecture was observed with and without nucleotide, our data suggest that binding to dsRNA induces the cleavage-competent state, whereas ATP acts later to remodel the complex and promote cleavage.

The stacked DRH-1 helicases are reminiscent of the filamentous assemblies formed by the RLR MDA5. However, we found that DRH-1 uses a distinct set of interactions to facilitate stacking interactions. The presence of nucleotide differentially redistributed helicase–helicase and helicase–RNA contacts in the AVC, as for MDA5, albeit the specific interactions and remodeling differed. Changes correlated with increased inter-helicase tilt and major groove widening in the dsRNA at helicase interfaces. Together, these findings provide a structural framework for understanding how DCR-1, DRH-1, and RDE-4 cooperate to promote viral dsRNA cleavage.

## RESULTS

### Overall structure of DCR-1•DRH-1•RDE-4 in a cleavage-competent state

Our previous work showed that dsRNAs of 42 or 52 basepairs bind to the AVC either by interacting directly with DCR-1 or with the helicase domain of DRH-1. In both cases, dsRNAs were not positioned within the catalytic RNase III (RIII) domains of DCR-1, suggesting the structures represent initial substrate-binding states^20^. In these studies, datasets containing no nucleotide or ATPγS resulted in the same 3D reconstruction, which allowed us to combine those datasets for processing. Despite containing a mixed nucleotide state, for simplicity, we refer to these structures as apo_initial_. To determine how the AVC transitions into a cleavage-competent state, we assembled the complex with dsRNA comprised of two 106nt strands that anneal to generate 2nt 3’ overhangs (ovrs) at both termini (hereafter called 3’ovr 106-dsRNA) and incubated the sample with ATP at 20°C prior to cryo-EM grid preparation. Under these conditions, we obtained a 3.0Å resolution 3D reconstruction of the complex in a cleavage-competent conformation (Figures 1 and S1). Notably, the reconstruction revealed dsRNA threaded through three stacked helicase domains: that of DCR-1, and two copies of DRH-1.

The density and model are oriented with DCR-1 stacked above two DRH-1 molecules (Figures 1B and 1C). We also observed density corresponding to one RDE-4 dsRBM interacting with the Hel2i subdomain of the DCR-1 helicase (Figures 1B and 1C). RDE-4 contains three structurally homologous dsRBMs^21,22^, and AlphaFold3 predictions support the C-terminal dsRBM (dsRBM3) of RDE-4 interacting with DCR-1’s Hel2i (Figures S2A and S2B). This is consistent with interactions of dsRBMs in accessory proteins included in structures of Dicer from other organisms^23–25^. In our model, a β-strand (β3) and an α-helix (α2) of the αβββα dsRBM fold contact α-helices 1 and 4 of Hel2i (Figures 1C, S2A, and S2C), similar to the interactions observed in structures of human DCR-1•TRBP and Drosophila Dcr2•R2D2^23–25^.

The DRH-1 molecule directly beneath DCR-1 displayed well-resolved density for the full-length protein, including its N-terminal domain (NTD), providing the first high-resolution structural information for this region. Directly below DRH-1 was another DRH-1 consisting only of its helicase and CTD domains (designated as DRH-1* to distinguish it from the DRH-1 directly contacting DCR-1) (Figures 1B and 1C). It is likely that the NTD of DRH-1* and adjacent regions of its assembly were too flexible to resolve. This is consistent with our previous work in which only the helicase domain and CTD of DRH-1 were resolved bound to dsRNA, despite having applied the entire complex to grids^20^.

Continuous density for dsRNA was threaded through the helicase domains of DCR-1 and both DRH-1 molecules (Figure 1B). Although the complex was assembled with a 3’ovr 106-dsRNA, we were only able to model ∼82nts of each strand within the reconstruction (Figure 1C). This missing ∼24nts is consistent with a single ATP-dependent cleavage event occurring during sample preparation, in agreement with our biochemical data showing that initial cleavage of a 2nt 3′ovr 106-dsRNA produces a 24nt siRNA product^20^. Thus, our structure likely represents the AVC after one round of endonucleolytic dsRNA cleavage, with the substrate poised for subsequent processing events.

### DCR-1 adopts a conserved cleavage-competent state

Our structure revealed that DCR-1 engages dsRNA such that it is positioned within the catalytic center formed by the intramolecular dimerization of RIIIa and RIIIb (Figures 2A–2C). Consistent with other Dicer structures, the RIII domains contact the sugar-phosphate backbone of both dsRNA strands with two negatively charged pockets containing the conserved catalytic residues E1416, D1420, D1575, E1578 (RIIIa) and E1682, D1686, D1791, E1794 (RIIIb) (Figures 2A–2D and S3A). In bacterial and yeast RNase III enzymes, these residues coordinate two Mg^2+^ ions, MgA and MgB, and two water molecules per ion^26,27^. MgA lowers the pKa of a bound water, enabling it to serve as the nucleophile for direct attack at the gamma phosphate, while MgB stabilizes the leaving group by neutralizing the developing negative charge on the 3′ oxygen^26,27^.

**Figure 2.**
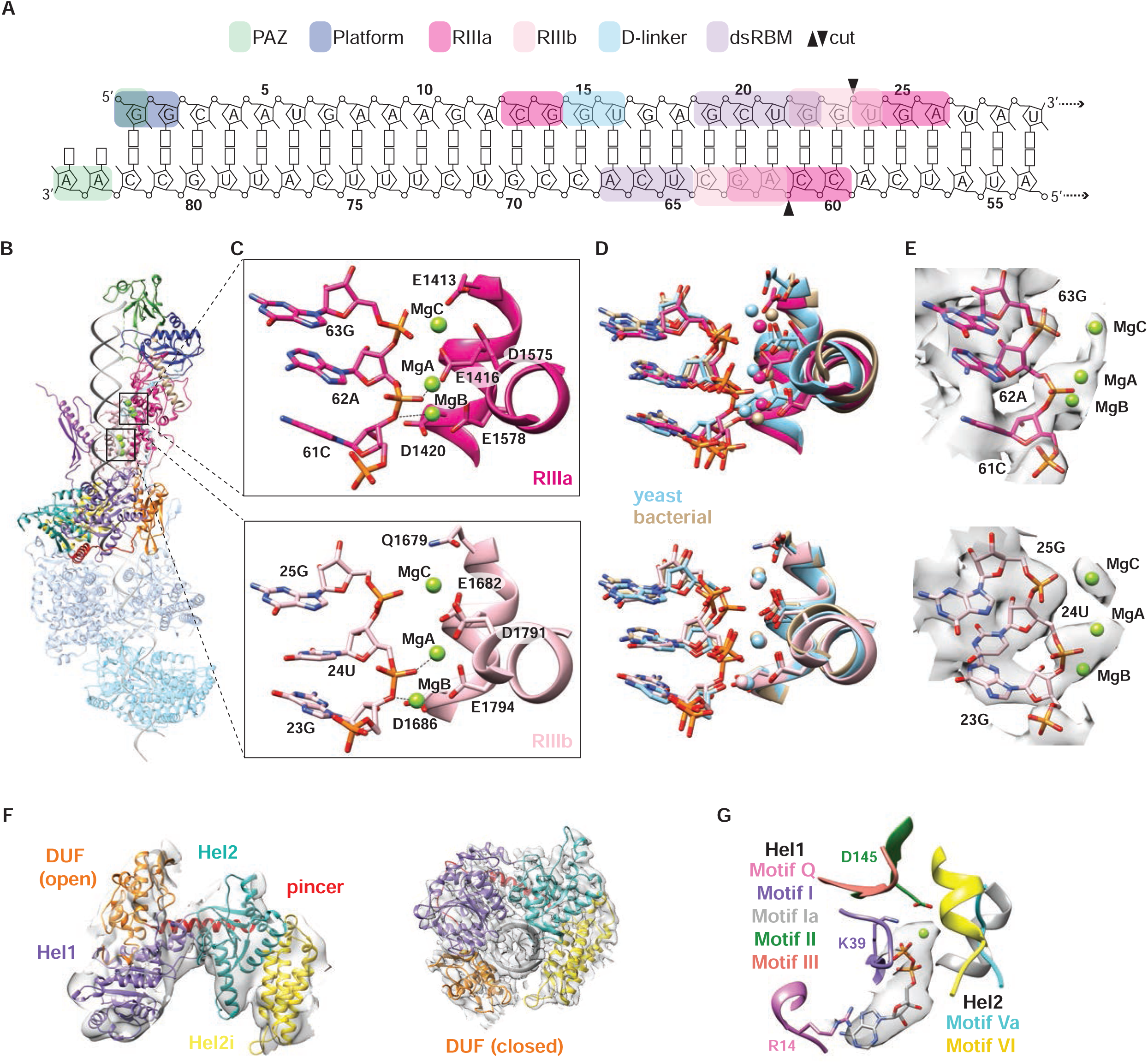
*C. elegans* DCR-1 adopts a conserved cleavage-competent state. A. Protein–RNA interactions of DCR-1 in a cleavage-competent state. Domains are colored as indicated, and black arrowheads indicate cleavage sites. B. Overview of DCR-1 colored by domain as in Figure 1A, except for DRH-1 and DRH-1* which are colored by protein as in Figure 1B and made transparent. RDE-4 is not visible in this view. C. Close up of DCR-1’s catalytic sites in the RIIIa (top) and RIIIb (bottom) domains. Mg^2+^ ions are represented as green spheres, and dashed lines indicate Mg^2+^ ion interactions proposed to facilitate cleavage of the phosphate backbone^26,27^. D. Magnified view of the structural alignment between *C. elegans* RIIIa (dark pink) and RIIIb (light pink) with bacterial RNase III (2NUG, tan)^26^ and yeast Rntp1 (4OOG, blue)^27^. E. Magnified view of density map overlayed with indicated nucleotides and Mg^+2^ ions. F. An AlphaFold model of DCR-1 helicase domain in an open conformation is overlayed with our previously obtained density (left, EMD-43430)^20^ and the closed conformation of our model is overlayed with our new density (right) (Supplemental Video 1). G. ADP and Mg^2+^ are overlayed with the reconstruction density and surrounding helicase motifs in DCR-1 are indicated.

Although only these two Mg^2+^ ions are required for cleavage, bacterial and yeast RNase III structures contain additional Mg^2+^ ions near the catalytic site^26,27^. Structural alignment of our reconstruction to the bacterial and yeast RNase III structures revealed extra density that overlaid well with a third Mg^2+^ ion, MgC, in both catalytic pockets (Figures 2C–2E). The proximity of polar residues such as E1413 in RIIIa and Q1679 in RIIIb suggested these residues stabilize MgC (Figure 2C). Our model explains how RIIIa cleaves between 61C and 62A, while RIIIb cleaves between 23G and 24U (Figures 2A and 2C). This cleavage reaction would generate a 23nt siRNA product that contains the canonical 2nt 3′ overhangs. Our previous biochemical studies showed that the first cleavage of a 3’ovr 106-dsRNA produces a 24nt siRNA product, while subsequent cleavage events yield 22nt or 23nt siRNAs^20^. Hence, the cleavage site in our model is consistent with the complex having completed one round of cleavage and poised for a second.

Surrounding the negatively charged RIII pockets are positively charged surfaces important for binding dsRNA, including the PAZ domain and dsRBM of DCR-1 (Figure S3B). Although these regions displayed poorer local resolution, their quality was sufficient to model them in agreement with other Dicer structures (Figures S3C and S3D). We modeled a 2nt 3′ovr extending into the PAZ domain (Figure S3E), consistent with the expected geometry of Dicer products and as reported for Dicers from other organisms^3^.

In one of our previously reported apo_initial_ structures of the AVC, dsRNA was bound to DRH-1, but not DCR-1^20^. In this structure, the helicase domain of DCR-1 was in an open conformation in which its DUF domain was positioned behind the Hel1 domain (Figure 2F, left, and Supplemental Video 1). In contrast, our cleavage-competent structure captured DCR-1’s helicase domain in a closed conformation, with its DUF domain repositioned in front of Hel1, allowing the helicase to enclose dsRNA (Figure 2F, right, and Supplemental Video 1). The helicase domain of dmDcr-2 also toggles between open and closed conformations, with the closed conformation associated with ATP-dependent dsRNA threading and processive cleavage^28,29^.

In our cleavage-competent structure, we modeled ADP within the conserved nucleotide-binding pocket of the DCR-1 helicase, formed by motifs from Hel1 and Hel2 (Figure 2G). ADP was also present in the ATP-binding pockets of both DRH-1 helicases (Figure S4). In all three sites, critical residues important for ATP specificity (DCR-1: R14, DRH-1: R294), ATP binding (DCR-1: K39, DRH-1: K320), and ATP hydrolysis (DCR-1: D145, DRH-1: D430), were positioned appropriately for catalysis (Figures 2G and S4). Our recent phylogenetic analyses indicate that the nematode ancestor of Dicer’s helicase domain retained ATP hydrolysis, but lost its ability to translocate along dsRNA, with DRH-1 providing this activity in the extant *C. elegans* AVC^30^. Consistent with this study, our structure shows that *C. elegans* DCR-1 retains the ability to adopt a closed conformation and hydrolyze ATP, whereas DRH-1 is well positioned to provide the translocation activity required for viral dsRNA processing.

### Progression to the cleavage-competent state does not require ATP

To determine whether ATP is required for formation of the cleavage-competent state, we collected another cryo-EM dataset of the AVC under the same conditions, but without ATP (Figures 3 and S5). Unexpectedly, this reconstruction closely resembled our cleavage-competent structure containing ATP, with the helicase domain of DCR-1 stacked above the helicase domains of two DRH-1 molecules (Figures 1 and 3A). Structural alignment revealed low overall RMSD values consistent with strong global similarity between the two models (Figure S6). As expected, this apo cleavage-competent structure (apo_CC_) lacked densities corresponding to nucleotide in any of the helicases (Figure 3B). However, we also observed several additional features that were only present in our apo_CC_ dataset.

**Figure 3.**
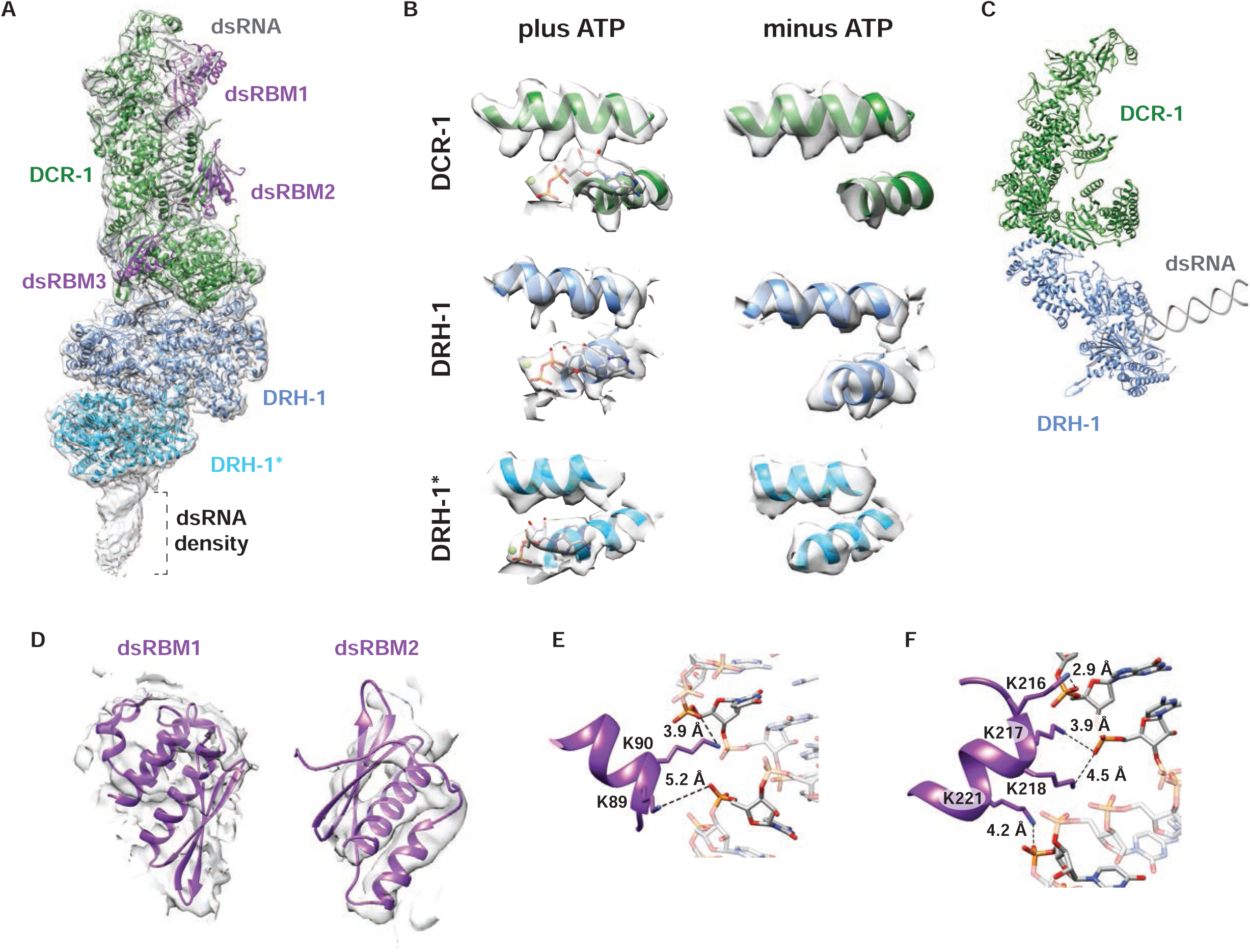
Structure of the AVC without ATP. A. 3D reconstruction of the AVC with 106nt 3’ovr dsRNA and without ATP. The model of each protein is fit into the density as indicated. RDE-4 is shown in purple with each dsRBM indicated. Additional density for dsRNA is indicated. B. Density of ATP-binding pocket for plus and minus ATP models shown for each helicase. All densities are displayed at the same threshold. C. Model of the complex in the initial binding state was built using EMD-43430 density^20^ and AlphaFold models of *C. elegans* DCR-1 and DRH-1. 52 blunt dsRNA was built in Chimera. D. Model of dsRBM1 and dsRBM2 overlayed with density. E. Interactions between RDE-4’s dsRBM1 with dsRNA. F. Interactions between RDE-4’s dsRBM2 with dsRNA.

In the apo_CC_ AVC, additional dsRNA density extended through the bottom of DRH-1* (Figures 3A and S6), consistent with an uncleaved, full-length 3’ovr 106-dsRNA. However, due to limited local resolution, we did not model this additional RNA density explicitly (Figure S6, see Methods). Additionally, a small subset of 2D classes from the apo_CC_ dataset showed the AVC in the initial binding state we previously reported, where DRH-1’s helicase domain is positioned away from DCR-1 (Figures 3C and S5B, red box)^20^. This initial binding state was not observed in our cleavage-competent ATP-bound dataset (Figure S1B). Taken together, these observations suggest that binding to the uncleaved 3′ovr 106-dsRNA is sufficient to promote the cleavage-competent conformation, while ATP binding and hydrolysis drives subsequent cleavage.

### RDE-4 dsRBM2 positions the dsRNA into DCR-1’s active site

Our apo_CC_ reconstruction also revealed additional densities that correspond to the first two dsRBMs of RDE-4 (dsRBM1 and dsRBM2), which were not observed in our other structures (Figures 3A and 3D). Although dsRBM1 had poorer local resolution than dsRBM2, both motifs were included in our model because their placement is consistent with the structure of dmDcr2•R2D2^23^ and supported by AlphaFold predictions (Figure S7). In both cases, dsRBM1 and dsRBM2 adopt similar orientations to those in our model.

Our model showed dsRBM1 making minimal contact with the dsRNA, with only a single lysine residue engaging the phosphate backbone (Figure 3E). In contrast, dsRBM2 contained a cluster of lysine residues positioned for contacting the phosphate backbone. Notably, K217 and K218 are within this cluster of lysine residues (Figure 3F), and a previous study showed that substitution of these residues with alanine impairs RDE-4 binding to dsRNA^31^. The limited contacts between dsRBM1 and dsRNA support previous biochemical studies showing that dsRBM1 is dispensable for promoting Dicer cleavage activity^22^, but is required for siRNA binding, where it is proposed to mediate transfer of siRNAs to Argonautes for downstream RNAi steps^21^. Our model further supports dsRBM2 as the most important dsRNA-binding motif for RDE-4^21,22^. Given that we only saw density for dsRBM2 in our cleavage-competent structures when ATP was absent, we favor a model in which dsRBM2 functions in apo_cc_ to position dsRNA into DCR-1’s active site as the complex transitions to the cleavage-competent state.

### DRH-1 NTD contains unique tandem CARDs

Our structure revealed that the NTD of DRH-1 is composed of two tandem helical bundles that resemble the CARD1 and CARD2 found in other RLR proteins (Figure 4). When aligned separately, each helical bundle superimposed well with the corresponding CARD domain of human RIG-I (Figures 4B and 4C). The tandem CARDs of the RLRs RIG-I and MDA5 are highly similar in both structure and function, as they interact with the same adaptor protein to stimulate the interferon response^32–34^. We aligned the CARDs of RIG-I and MDA5 (Figure S8A) to illustrate that our comparison of the CARDs between DRH-1 and RIG-I is similar to superimposing those domains with MDA5.

**Figure 4.**
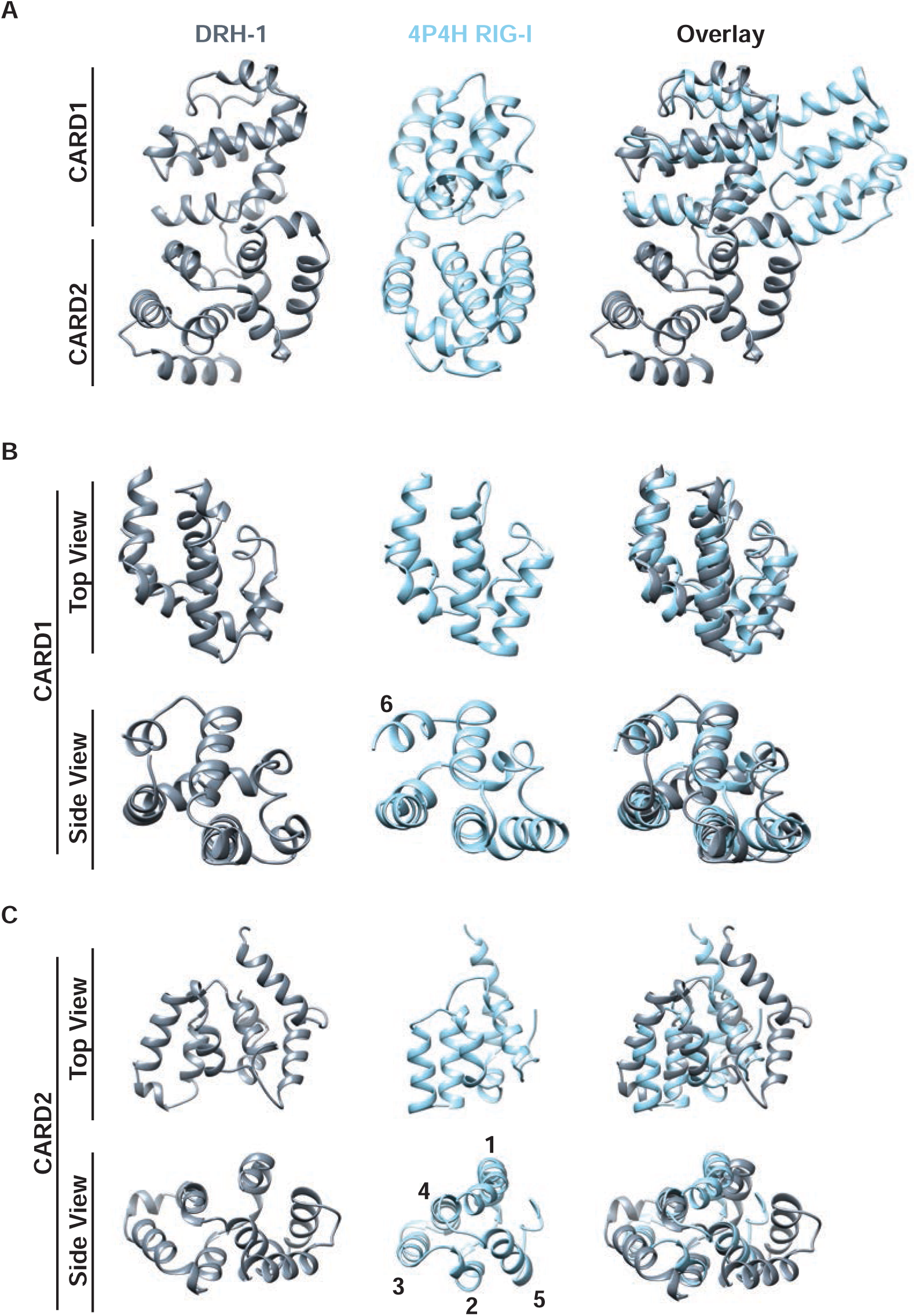
DRH-1 NTD contains unique tandem CARDs. A. Structure of the DRH-1 NTD (left), RIG-I CARDs (4P4H, middle)^34^, and overlay between both (right). Structures are oriented to show CARD1 above and CARD2 below. B. Top and side views of CARD1 from DRH-1 (left) and RIG-I (4P4H, middle)^34^, and an overlay between both. C. Top and side views of CARD2 from DRH-1 (left) and RIG-I (4P4H, middle)^34^, and an overlay between both. Helices 1 – 5 in RIG-I are indicated.

*C. elegans* encodes another RLR homolog known as DRH-3 that also contains two CARD-like α-helical bundles^35^. However, structural comparison revealed that the tandem CARDs of DRH-1 and DRH-3 are much less similar (rmsd values > 5Å) (Figures S8B–S8D). Previous studies show DRH-1, but not DRH-3, stimulates the intracellular pathogen response (IPR)^36,37^. Furthermore, another study shows that although RIG-I’s helicase and CTD domains can be swapped with the analogous domains of DRH-1 and still activate an antiviral response in *C. elegans*, the CARD domains were not interchangeable^18^. Thus, DRH-1 contains unique tandem CARDs that are functionally distinct from those of RIG-I and MDA5. This specialization likely underlies the role of DRH-1 in initiating the IPR in *C. elegans*, rather than the canonical interferon pathway.

### The CARD-Helicase Linker of DRH-1 mediates the DCR-1•DRH-1 interaction

Our previous low-resolution apo_initial_ AVC structures suggested that the NTD of DRH-1 mediates contact with DCR-1, but the interaction interface could not be defined. Our cleavage-competent structures enabled detailed modeling of DRH-1’s interaction with DCR-1, where we discovered the interaction was not mediated by the NTD and instead a linker adjacent to the NTD. RLRs contain a linker region that separates the N-terminal CARDs from their helicase-CTD RNA binding module (CHL, CARDs-Helicase Linker)^38^. DRH-1 contains an analogous ∼67 amino acid linker, which we also refer to as CHL (Figure S9A). Unlike the flexible linkers in other RLRs, the CHL of DRH-1 is resolvable, including an α-helix and a two-stranded β-sheet (Figure S9). No other RLR contains a modeled linker due to the apparent intrinsic flexibility of this region. Consistent with our structure, AlphaFold predicts that the DRH-1 CHL is partially structured, whereas the corresponding linkers in other RLRs are predicted to be unstructured (Figure S9).

Our cleavage-competent structures revealed that the CHL forms a key point of contact between DCR-1 and DRH-1 by extending a β-sheet with the Hel2 domain of DCR-1 (Figures 5A–5C). Given that the pincer domain is important for stabilizing MDA5 oligomerization into filaments^39^, we were intrigued to see DCR-1’s pincer domain near the CHL (Figure 5B). However, upon close inspection, the pincer is positioned too far away to contribute directly to the interface between DCR-1 and DRH-1. A second interface is formed between the DUF domain of DCR-1 and the CTD of DRH-1 (Figures 5A, 5D, and 5E). This interaction was not observed in our previous apo_initial_ state (Figures 3C and 5F). Comparison of these structures indicates that when the complex enters the cleavage-competent state, DCR-1’s helicase closes (Figure 2F) and stacks with DRH-1’s helicase, bringing the DUF and CTD into contact (Figures 5A, 5D, and 5E). We therefore propose that the CHL serves as a structural tether that maintains the interaction of DCR-1 and DRH-1 during the large conformational rearrangements between the two states. This interaction may position DRH-1 so that it does not sterically hinder DCR-1 from toggling between its open and closed conformations (Supplemental Video 2).

**Figure 5.**
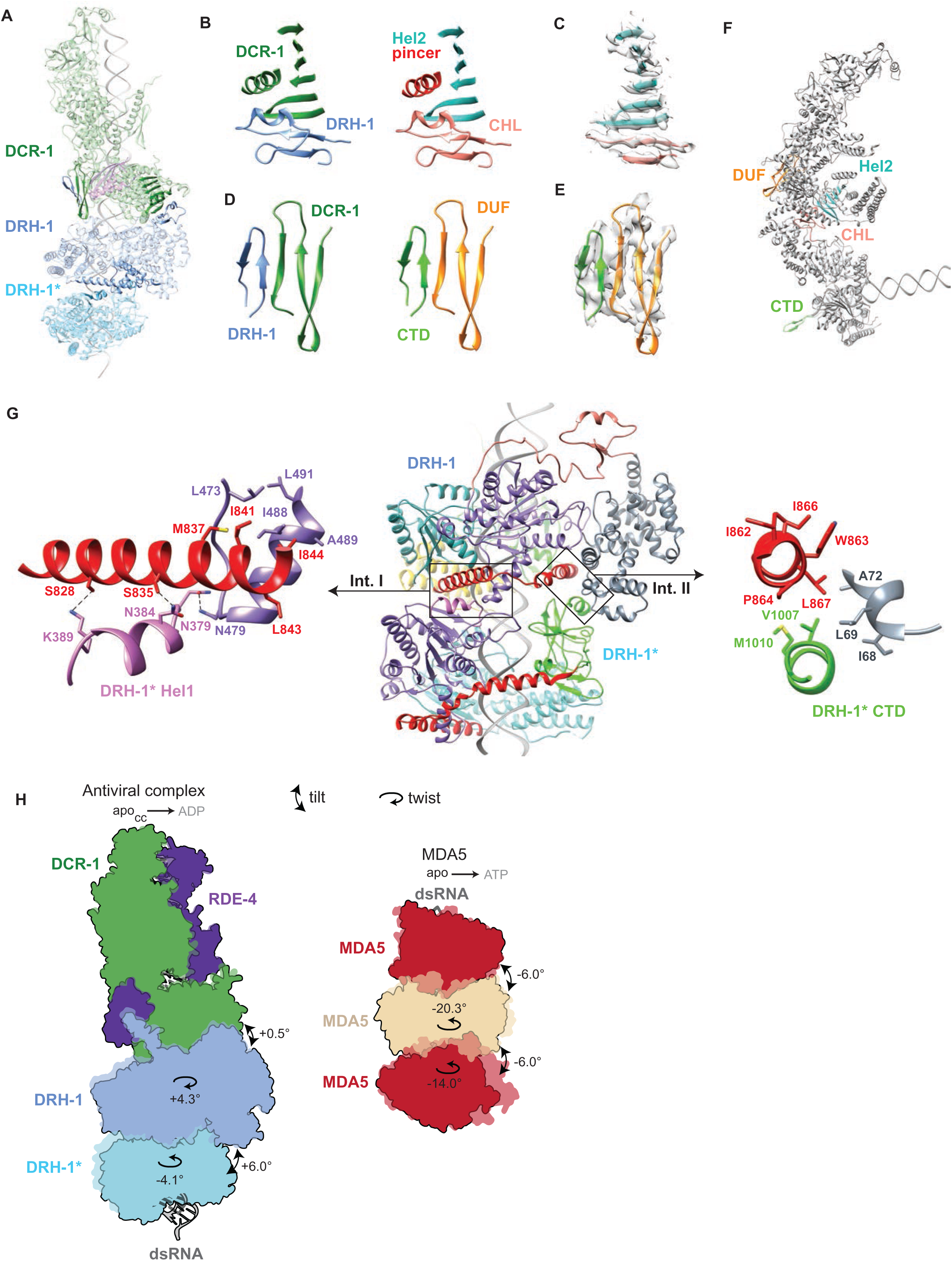
Interactions between DCR-1•DRH-1 and DRH-1•DRH-1. A. Overview of the interactions observed between DCR-1•DRH-1 and DRH-1•DRH-1*. The interactions are displayed as solid whereas the rest of the complex is transparent. B. Magnified view of contact between DCR-1 and DRH-1. On the left, proteins are colored as indicated in (A) and on the right domains are colored as indicated. C. Density of DCR-1’s Hel2 forming an extended β-sheet with DRH-1’s CHL. D. Magnified view of contact between DCR-1 and DRH-1. On the left, proteins are colored as indicated in (A) and on the right domains are colored as indicated. E. Density of DCR-1’s DUF forming an extended β-sheet with DRH-1’s CTD. F. Model of the complex in the initial binding state with contacts colored as in B and D. G. Magnified view of interactions facilitating DRH-1•DRH-1* stacking. Left, pincer and Hel1 domains of DRH-1 colored as indicated; DRH-1* Hel1 colored in pink. Right, DRH-1 pincer and NTD domains colored as indicated; DRH-1* CTD colored in green. Locations of these interactions in the protein complex are in (A), where relevant regions of DRH-1 and DRH-1* are displayed as solid. H. Two-dimensional schematics of the AVC and MDA5 trimer illustrating structural changes between nucleotide states. The apo (outlined) and ADP or ATP-bound (transparent, no outline) models are overlaid for both the AVC and MDA5. Changes in twist and tilt between adjacent helicases are indicated. The method used to measure twist and tilt is described in Figure S11.

### DRH-1 forms oligomers mediated by electrostatic and hydrophobic interactions

Our structure revealed that DRH-1 forms dimers in the presence of DCR-1, providing the first observation of DRH-1 oligomerization. This arrangement is reminiscent of the filamentous assemblies formed by its ortholog MDA5. MDA5 forms head-to-tail filaments mediated by interactions of the pincer domain of one MDA5 monomer and the Hel1 and CTD domains of another (Figure S10). DRH-1 uses these same domains to form its dimer interface, but the underlying interactions differ. Whereas the hydrophobic residues in MDA5’s pincer domain form intermolecular interactions, the corresponding residues in DRH-1 instead stabilize through intramolecular packing (Figures 5G and S10).

The intermolecular interactions in interface I (Int. I) of the DRH-1 dimer are mediated by the N-terminal α-helix of DRH-1’s pincer domain contacting Hel1 of DRH-1* (Figure 5G, Int. I). In this interface, S828 and S835 form polar interactions with Hel1 of DRH-1*, while an additional interaction is formed between N479 of DRH-1 and N379 of DRH-1*. These interactions are reinforced by an intramolecular hydrophobic pocket comprised of residues in Hel1 (L473, I488, A489, L491) and pincer (M837, I841, L843, I844).

Interface II (Int. II) also differs from that of MDA5. In MDA5, this interface is mediated by the pincer domain of one monomer and the CTD of the next monomer (Figure S10). In contrast, in Int. II of DRH-1, the C-terminal α-helix of DRH-1’s pincer packs against the NTD of the same DRH-1 molecule through hydrophobic interactions, and this interaction is further stabilized by hydrophobic contacts with the CTD of DRH-1* (Figure 5G, Int. II). Interestingly, a mutation in DRH-1’s NTD (A72T) maps to this region and reduces antiviral RNAi activity in *C. elegans*^40^. This suggests that, like MDA5, disrupting DRH-1 oligomerization may compromise function.

### Nucleotide binding remodels inter-helicase twist and tilt

Although the apo_CC_ and ADP-bound AVC models were globally similar (Figure S6), closer inspection revealed nucleotide-dependent differences in the relative orientations of the stacked helicases. We quantified changes by measuring twist and tilt between adjacent helicase domains (Figure S11). Twist was defined as the angular displacement between pincer-domain centroids, and tilt was calculated as the angle between normals of the best-fit planes of the helicase cores. Because the dsRNA deviates from perfect linearity (see next section), the calculated helical axis, and thus twist measurements, are approximate.

Transition from the apo_CC_ to the ADP-bound state resulted in a +0.5° increase in tilt between DCR-1 and DRH-1, and a +6.0° increase between DRH-1 and DRH-1* (Figure 5H). Twist changes were also interface-specific: twist increased by +4.3° between DCR-1 and DRH-1 but decreased by −4.1° between DRH-1 and DRH-1*. Thus, nucleotide binding correlated with increased tilt at both interfaces, whereas twist changes occurred in opposite directions across adjacent helicase pairs.

Despite these conformational shifts, the interface interactions between DCR-1 and DRH-1 remained largely unchanged (Figures 5B–5E), showing only minor increases in buried surface area when transitioning from apo_CC_ to ADP-bound state (<3%; Figure 5B interactions +59Å², Figure 5D interactions +17Å², respectively). Contacts with dsRNA were also relatively unchanged (<1% decrease) for both DCR-1 and DRH-1 (−13.5 and −14Å², respectively). In contrast, contacts between the two DRH-1 helicases were substantially weakened. Interface I (Figure 5G) decreased by a modest 3% (−19Å²), but interface II decreased by 26% (−152Å²), upon transitioning from the apo_CC_ to ADP-bound state. Additionally, a small contact between Hel2i of DRH-1 and Hel2 of DRH-1* (171Å²) was completely lost as the two DRH-1 molecules tilt away from one another (Figure 5H).

Notably, weakening of the inter-helicase contacts between DRH-1 and DRH-1* coincided with an ∼4% increase in engagement of DRH-1* with dsRNA (+105.5Å²). Thus, nucleotide binding in the AVC slightly strengthens the DCR-1–DRH-1 interface while loosening the DRH-1– DRH-1* interface and concurrently enhancing RNA engagement by the terminal DRH-1* helicase.

Comparing this to MDA5, prior work from the Modis lab^39^ described nucleotide-dependent changes in helicase twist and accompanying tilt changes, which we quantified here using our method as a uniform −6.0° decrease in tilt at each helicase interface, when transitioning from the apo to the ATP-bound state, and decreases in twist of −20.3° and −14° for the middle and bottom helicases relative to the top (Figure 5H). In contrast to the AVC, both interfaces in MDA5 undergo coordinated decreases in twist during this transition. MDA5 has three regions of contact between adjacent helicases that are equivalent at each interface; therefore, only the middle-bottom pair is considered. Upon nucleotide binding, interface I (Figure S10) as well as interface II (Figure S10) both decrease in buried surface area by 20% (−88 and −72Å, respectively). A small interface between Hel2i and Hel2, similar to the contact between the same domains in the AVC’s DRH-1 domains, increases by 66% (+107Å²) as the helicases change tilt relative to each other (Figure 5H). Overall, these changes result in a net decrease in inter-helicase contact, in a manner similar to the DRH-1-DRH-1* interface in the AVC, albeit to a lesser extent. This weakening of helicase-helicase interactions for MDA5 is accompanied by a substantial increase in helicase-dsRNA contact of 16% (+379Å²). Thus, nucleotide binding weakens helicase packing, but increases RNA engagement for both MDA5 and DRH-1, but these changes are associated with decreases in tilt/twist for MDA5 and increases in tilt/twist at the DRH-1-DRH-1* interface in the AVC.

### RNase III and helicase binding distorts the dsRNA from A-form geometry

Another striking feature in our structure was the substantial distortion of the dsRNA away from canonical A-form geometry (Figure 6A). Using CURVES+^41^, we identified regions where the major groove width exceeded 10Å, approximately double the ∼4.9Å expected for ideal A-form dsRNA (Figure 6B).

**Figure 6.**
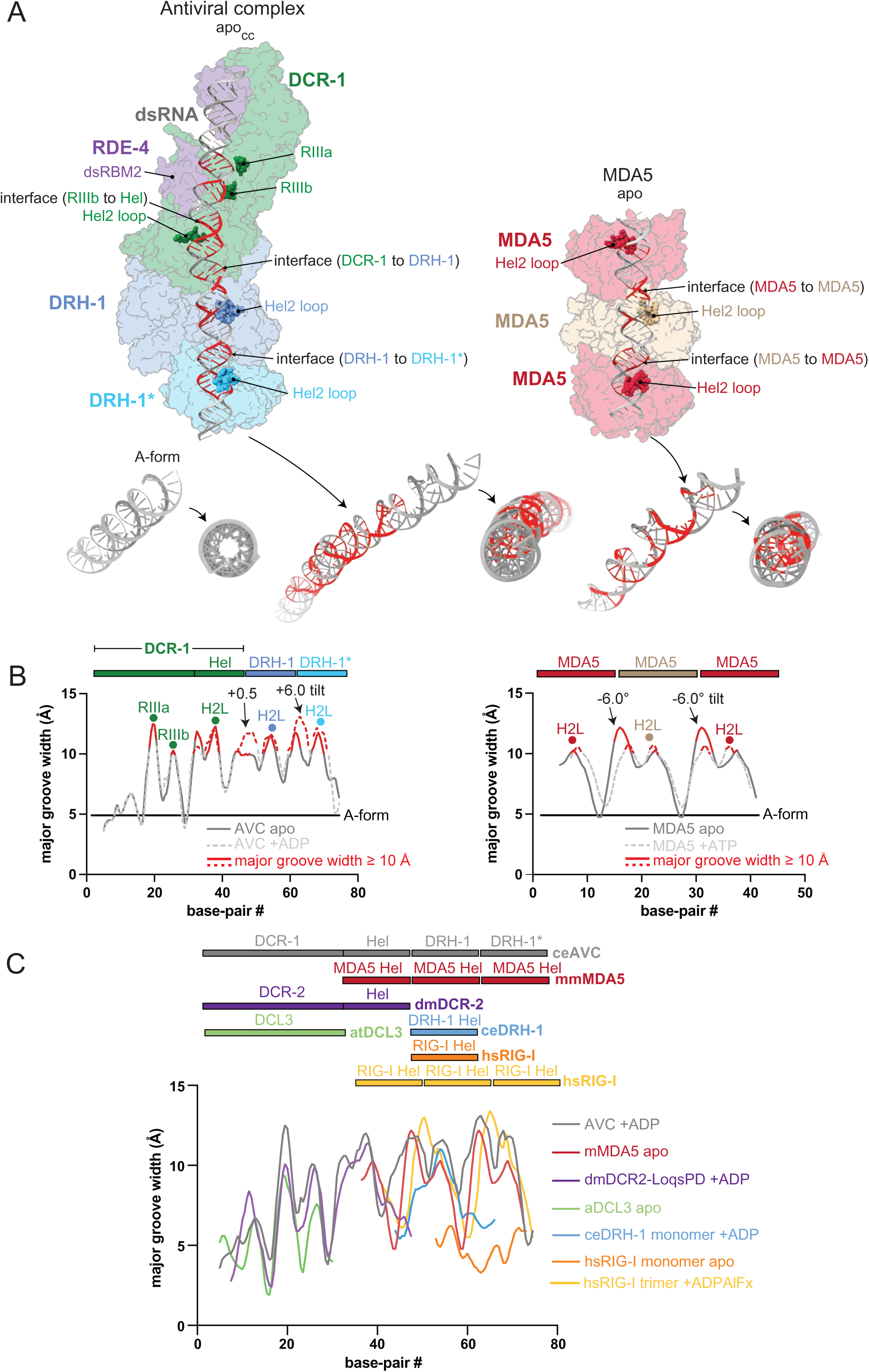
Antiviral complex and MDA5 binding distort dsRNA away from canonical A-form geometry. A. The apo antiviral complex (left) and MDA5 (right) models are shown^33,39^. Major groove widths above ≥10Å, as defined by CURVES+^41^, are colored red and correspond to Hel2 loop insertions and the RNase IIIa/b binding sites as well as to the interface between helicases or between Dicer’s helicase and the RNAse IIIb domain. The RNase IIIa/b catalytic residues and Hel2 loops for DCR-1, DRH-1, and MDA5 are highlighted using a space-filling model. Shown below the models are the respective bound dsRNAs at a diagonal and helical views to illustrate the degree of disruption from a perfect A-form helix, which is depicted on the left as a comparison. B. Graphs of major groove widths for A-form dsRNA and the dsRNA associated with the antiviral complex and MDA5 trimer at different nucleotide bound states. Solid and dotted grey lines lines show apo and ATP-bound models, respectively. In the AVC, the apo_CC_-to-ADP transition increases the DCR-1-DRH-1 helicase interface groove width by 0.5°, and the DRH-1-DRH-1* width by 6°. In MDA5, the apo-to-ATP transition decreases both interface groove widths by 6°. The lines are colored red in the same manner as in (A), and the locations of the RNase IIIa/b catalytic residues, Hel2 loops, and interfaces are indicated as dots above the lines. The location of the domains of the antiviral complex and MDA5 trimer relative to the dsRNA is indicated above the plots. C. Graphs of major groove widths are shown for the dsRNA in structures of the apo_CC_ AVC (grey), mouse no nucleotide MDA5 trimer (red)^33,39^, ADP bound Drosophila melanogaster Dicer-2-Loqs-PD complex (dmDCR2-LoqsPD, purple, 7W0E)^28^, apo Arabidopsis Dicer-Like Protein 3 (aDCL3, green, 7VG3)^56^, *Caenorhabditis elegans* ADP-bound DRH-1 monomer (ceDRH-1, blue, 8T5S)^20^, apo human RIG-I monomer (dark orange, 8SCZ), and ADPAlFx-bound RIG-I trimer (light orange, 7JL3)^45^. mMDA5, ceDRH-1, and hRIG-I monomer models were aligned by superimposing the basepairs contacted their Hel2 loops onto the corresponding basepairs contacted by DRH-1’s Hel2 loop in the antiviral complex model. dmDCR-2 and aDCL3 were aligned using the basepairs contacted by the RNase IIIa/b catalytic residues. The domain structure of the different proteins/complexes relative to the dsRNA is indicated above the plots.

As illustrated, each of the three helicases in the AVC correlated with two peaks of major groove widening (Figures 6A and 6B, red). One correlated with an insertion of a loop of the Hel2 subdomain (Hel2 loop) into the major groove (Hel2 loop, residues 756-767 for MDA5, 423-437 for DCR-1, and 732-750 for DRH-1), while a second widening occurred at dsRNA positioned at the interfaces between the stacked helicases, or between DCR-1’s helicase and RNase IIIb domain. This pattern resembles the major groove widening caused by Hel2 loop insertion observed within the MDA5 monomer^33,39^, and we likewise detected groove expansion at Hel2 loop insertion sites and interfaces between stacked helicases in the trimeric model of MDA5 (Figures 6A and 6B).

Another cluster of major groove widening was observed in the AVC at the RNase III catalytic residues and overlapped with the binding site of RDE-4 dsRBM2. Similar distortions have been reported in crystal structures of bacterial and yeast RNase III enzymes, although the widening in these structures is less pronounced^27,42,43^.

The regions of major groove widening also coincided with increases in rise per bp, decreases in minor groove width, and increases in hydrogen-bond distances between basepairs for both MDA5 and DRH-1 (Figure S12). However, these changes were smaller in magnitude and more variable than the changes in major groove width, making major groove width the clearest indicator of dsRNA distortion in both systems.

While not previously reported for Dicers, nearly all deviations in major groove width we observed could be overlaid with those from previously solved structures, including mouse MDA5, the *Drosophila* Dicer-2-Loqs-PD complex, and *Arabidopsis* Dicer-Like 3, suggesting deformation of the bound dsRNA by these Dicer and related helicases is a conserved phenomenon (Figure 6C). Notably, we did not observe major groove widening in a structure of RIG-I bound to the dsRNA terminus, where its Hel2 loop acts as a gating mechanism to distinguish 5′-ppp from 5′-OH RNA^44^. However, widening reappeared in a separate structure in which three RIG-I molecules were bound internally along dsRNA (Figure 6C)^45^. The latter supports the conclusion that groove expansion in our structures arises specifically from Hel2 loop insertion into the major groove rather than being a general feature of helicase-RNA engagement.

## Discussion

Our prior Ancestral Protein Reconstruction (APR) analyses indicate the nematode ancestor of the helicase domain of extant *C. elegans* Dicer had minimal ATP hydrolysis, dsRNA binding, and translocation activity, and our biochemical analyses suggest *C. elegans* co-opted the dsRBP RDE-4 to resurrect dsRNA affinity, and the RLR DRH-1 to provide ATPase-dependent translocation activity^20,30^. Here we report cryo-EM structures of the AVC, in the absence and presence of ATP, providing insight into how DCR-1, DRH-1, and RDE-4 work together to resurrect the activities that in some invertebrates, such as dmDcr2, are intrinsic to a single protein^29^. A key feature of the cleavage-competent AVC structures reported here is the stacking of three helicase domains, that of DCR-1 with those from two molecules of DRH-1, reminiscent of the oligomerization that is observed with mammalian RLRs such as MDA5, LGP2 and RIG-I^39,45,46^.

### What is the initial step towards cleavage of viral dsRNA by the AVC?

While of lower resolution (6.1-7.6Å), our previously published structures of the AVC, performed without nucleotide, or with ATPγS, showed DRH-1 clamped around the end of the dsRNA in a closed conformation, not yet stacked on the helicase of DCR-1. As illustrated, we propose this represents an initial binding state (Figure 7, I DRH-1 initiation). In our prior studies we also observed a structure with dsRNA engaged as in I, with an additional dsRNA bound to DCR-1’s Platform•PAZ domain (Figure 7, II Platform•PAZ initiation). As discussed below, we speculate this structure signifies there is an additional way to enter the cleavage-competent state.

**Figure 7.**
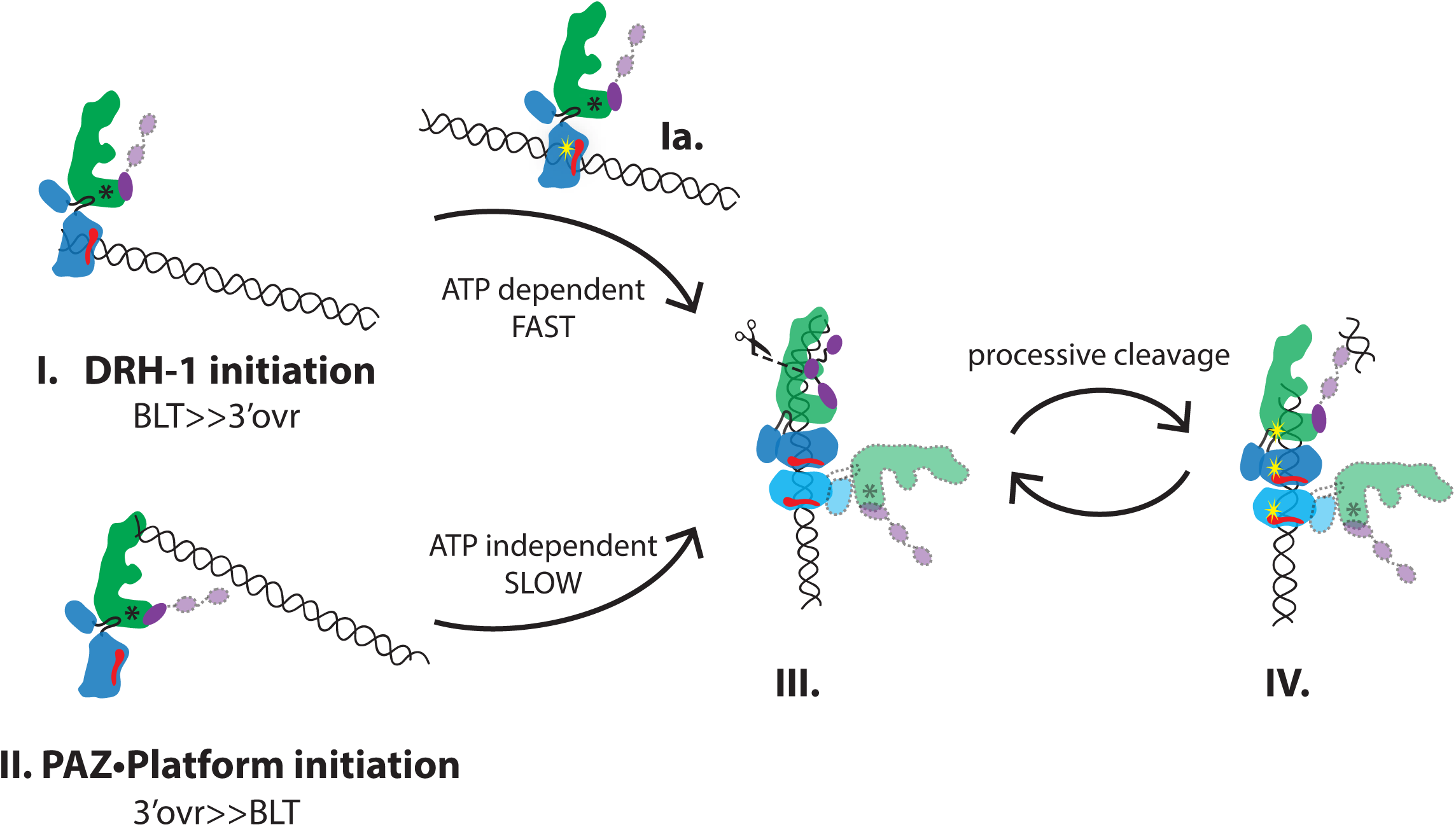
Model of the AVC entering the cleavage-competent state. Various intermediates lead to the cleavage-competent structures reported here, without (III) or with (IV) ATP; DCR-1 (green), DRH-1 (blue) and RDE-4 (purple). Existing data indicate blunt dsRNA prefers initiation by state I, and 3’ovr, state II. Once the AVC enters the cleavage-competent state (III), dsRBM2 of RDE-4 helps position dsRNA in the RNase III active sites (scissors) for cleavage. After the first cleavage, ATP-dependent translocation by DRH-1 repositions dsRNA for processive rounds of cleavage (step IV). siRNA products are passed to downstream components of RISC, including RDE-1, likely via interaction with dsRBM1. Red slash in DRH-1, pincer domain to illustrate helicase orientation. Black asterisks on DCR-1, OPEN conformation of helicase domain; absence of asterisk implies CLOSED conformation. Yellow stars, active ATP hydrolysis. Dotted lines mark domains whose presence are supported by previous biochemical data and low-resolution density in 2D classes^20^.

The apo_CC_ and ADP-bound cleavage-competent structures reported here both show an interaction of DRH-1’s CHL with DCR-1, and the higher resolution of these structures reveal that the nearby NTD, as for other RLRs, is comprised of two tandem CARDs, consistent with recent studies based on Alphafold models^37^. The higher resolution structures allowed us to model a specific interaction of DRH-1’s CHL, just C-terminal to the NTD, with the Hel2 domain of DCR-1 that extends a β-sheet (Figure 5B). Interestingly, in contrast to the CHL of mammalian RLRs, that of DRH-1 is resolvable and includes an α-helix and a two-stranded β-sheet (Figure S9).

Subsequent steps that allow the AVC to enter a cleavage-competent state require large conformational changes (Figure 7), and it is tempting to postulate that the interaction of DRH-1’s CHL serves to tether DRH-1 and DCR-1 during this transition. While this is our preferred model, the lower resolution of our prior structures makes it impossible to know if the interaction that occurs in the cleavage-competent state is the same in detail as that observed in the initial binding state.

As in other Dicer structures, dsRBM3 of RDE-4 was observed interacting with Hel2i of DCR-1 in both apo_CC_ and ADP-bound cleavage-competent structures. Only the apo_CC_ structure revealed all three dsRBMs (Figure 7, III.), and their positions were similar to dmDcr2•R2D2^23^, and also consistent with recent biochemical studies indicating dsRBM3 is important for tethering RDE-4 to DCR-1, dsRBM2 is important for binding dsRNA and as in other RNase III enzymes, likely for positioning dsRNA in the active site, with dsRBM1 well positioned for transferring the cleaved siRNA to a downstream Argonaute^22^. Possibly our ADP-bound state (Figure 7, IV.) represents a post-cleavage state where both dsRBM1 and dsRBM2 have disengaged as the cleaved siRNA is passed to the RISC.

### Progression to the cleavage competent state requires a large conformational change

In our model, as illustrated, either of the initial binding states can proceed to the cleavage-competent structure, which was observed in the apo_CC_ (III) and ADP-bound (IV) structures reported here (Figure 7). Progression to the cleavage-competent state requires a reorientation of DRH-1 to stack with DCR-1, stabilized by an interaction of DCR-1’s DUF with DRH-1’s CTD in addition to the interaction of the CHL with DCR-1’s Hel2 (Figure 5A). In both the apo_CC_ and ADP-containing structures an additional DRH-1 (DRH-1*) stacks below, reminiscent of the oligomerization of mammalian RLRs such as MDA5. While the domains that mediate stacking of MDA5, the pincer domain, Hel1 and the CTD, are also used for DRH-1 stacking, the specific interactions differ substantially (compare Figures 5G and S10).

### Integrating biochemical and structural observations: discrimination of self versus nonself termini

Our prior in vitro studies show remarkably similar biochemical properties for the AVC, comprised of three proteins^20^, and dmDcr2, a single protein^47^. Both invertebrate antiviral activities cleave dsRNA in an ATP- and terminus-dependent manner, with the optimal reaction occurring with blunt dsRNA in the presence of ATP. For example, when comparing a 106-dsRNA in the presence of ATP, for both invertebrate activities blunt termini are cleaved far more efficiently than dsRNA with 3’ovr termini (*k*_obs_ min^-1^, AVC_blunt_, 0.14, dmDcr2_blunt_, 0.19; AVC_3’ovr_, 0.006, dmDcr2_3’ovr_, 0.01). As for studies of mammalian immune sensors, such as RIG-I, this terminus preference has been proposed to reflect a self versus nonself discrimination, based in the fact that dsRNA made by viruses often has blunt termini^48^. Structural and biochemical studies of dmDcr2 indicate the enhanced efficiency of blunt dsRNA cleavage, compared to that of 3’ovr dsRNA, arises because it is threaded through the helicase domain to allow processive cleavage^29^. The similarity of the AVC kinetic parameters suggest that blunt dsRNA also threads through DCR-1’s helicase domain within the AVC, and our structures showing dsRNA threaded through three stacked helicases are consistent with this model (Figures 1B and 7). Presumably because of the loss of the translocation activity in the nematode ancestor of Dicer’s helicase, DRH-1 molecules are stacked below DCR-1 to allow translocation along the dsRNA and to push the dsRNA into DCR-1’s helicase domain and the body of the enzyme. In our model we envision the threading of blunt dsRNA initiates from intermediate I. In our prior structures formed with blunt dsRNA, addition of ATP showed some 2D classes with DRH-1 localized internally on dsRNA (intermediate Ia) suggesting an ATP-dependent translocation step that may facilitate the conformational change that allows threading into DCR-1’s helicase domain.

However, and importantly, the AVC structures we report here were acquired with a 106- dsRNA with a 3’ovr. While transient kinetic assays indicate a small fraction of 3’ovr dsRNA can thread through dmDCR-1’s helicase domain, this terminus is predominantly cleaved in a non-processive, inefficient reaction that initiates with an interaction with the Platform•PAZ domain^20,49^. Consistent with this, while a small subset of 2D classes from the apo_CC_ structure showed the cleavage-<u>in</u>competent configuration of intermediate I (Figure S5), intermediate Ia was not observed even with the addition of ATP, suggesting translocation at this step is rarely accessed by a 3’ovr dsRNA. That said, the characteristic stacking of three helicases was observed in both the apo_CC_ or ADP-bound structures (Figure 7, III and IV), indicating that this structure does not require ATP-dependent translocation to form. Thus, in our model we show an alternate route to the 3-stacked helicase complex that initiates with an interaction of the 3’ovr with the Platform•PAZ domain (Figure 7, II). While a 3’ovr dsRNA strongly prefers to bind to the Platform•PAZ domain, our in vitro studies indicate cleavage of this substrate is inefficient and slow^20^, and possibly this enabled the capture of the cleavage-competent structures we report here. The rate limiting step of cleavage of a 3’ovr dsRNA by the AVC may involve the helicase domains of DCR-1 and DRH-1 closing around the dsRNA to form the stacking interactions observed in intermediate III. This may be a very slow step given that optimal interactions of the helicase domains require disruption of the A-form dsRNA structure, which we speculate would be far more optimal in the context of an ATP-dependent threading mechanism. However, once forced into the cleavage-competent state, in the presence of ATP (intermediate III), ATP hydrolysis likely promotes dsRNA translocation toward the RNase III domains. Regardless of whether intermediate III forms via threading from intermediate I, or from intermediate II by the dsRNA swinging in from the Platform•PAZ interaction to be clamped by the helicases, after the first cleavage to yield a 3’ovr, both complexes would be identical and be cleaved processively in the presence of ATP.

### The RIG-I like receptor family of helicases and their interaction with dsRNA

The narrow major groove of dsRNA makes it difficult for proteins to make sequence specific contacts, which largely occur in the major groove^50^. However, there are countless examples of major groove distortion that enable proteins and RNA molecules to make sequence-specific interactions^51–53^. All three helicases observed in our apo_CC_ and ADP-bound structures show two regions of major groove widening, one at each helicase-helicase interface and one within each helicase, involving Hel2 loop that is essential for function of mammalian RLRs^33,54,55^. In MDA5, deletion of the Hel2 loop has no impact on RNA binding but abolishes ATPase activity and prevents IFN signaling *in vivo*^33^. For RIG-I, phosphorylation of the Hel2 loop preserves RNA binding while reducing ATPase activity and impairing downstream signaling, with in vitro studies revealing defects in streptavidin displacement consistent with disrupted translocation^54,55^. These findings suggest that Hel2 loop-mediated deformation of the RNA is critical for productive ATPase cycling and signaling.

For the interfaces between MDA5 oligomers and DRH-1/DRH-1*, nucleotide binding weakens helicase packing but increases RNA engagement (Figures 5C and 6), albeit these changes are associated with decreases in tilt/twist for MDA5 and increases in tilt/twist at the DRH-1-DRH-1* interface of the AVC. Nucleotide-dependent remodeling of MDA5 is proposed to play a role in self-nonself dsRNA discrimination, and the changes in interface contact and RNA engagement are consistent with the proposed “testing” of the dsRNA’s stability^39^. As yet, the biological significance of the stacking of two DRH-1 molecules is unclear, but given the importance of DRH-1 for antiviral defense, possibly DRH-1 stacking similarly contributes to “testing” of dsRNA for properties of self versus nonself. In contrast to the changes that occur at the interfaces of these RLRs, the relatively minor changes in twist, tilt, and interactions between DCR-1 and DRH-1, and with dsRNA, upon nucleotide binding are likely related to the need to maintain a tight grip on the viral dsRNA as it is processively cleaved into siRNAs.

The second peak of major groove widening that we associate with the interface between stacked helicases is consistent with mechanical strain imposed on the dsRNA by inter-helicase contacts or neighboring domains (e.g., RNase IIIb), as an AVC structure in which only a single DRH-1 was observed bound to dsRNA lacks the second peak of major groove widening^20^ (Figure 6C). Supporting this, major groove widening at the helicase interfaces in both the AVC and MDA5 increase by ∼ 1.5Å (Figure 6B) with the increased tilt angle between stacked helicases across nucleotide states (Figure 5H). Possibly, this nucleotide-dependent increase in major groove widening may also contribute to self-nonself discrimination, as most endogenous dsRNAs have more mismatches and are edited by ADARs, and may be less able to withstand the strain applied by this widening, whereas perfectly base-paired viral dsRNAs could better accommodate it.

## Concluding Statement

Our AVC structures capture cleavage-competent states that reveal new roles for RLRs in antiviral defense. In vertebrates, RLRs recognize viral dsRNA to trigger an interferon response, whereas in the invertebrate *C. elegans*, the collaboration of the RLR DRH-1 with DCR-1 promotes antiviral RNAi. We believe the use of 3’ovr dsRNA, which is a suboptimal substrate of the AVC and cleaved very slowly, allowed the capture of the structures we report here. A key goal for future studies will be to determine a cleavage-competent structure with the optimal blunt dsRNA substrate.

## Acknowledgements

We acknowledge David Belnap and Barbie Ganser-Pornillos at the University of Utah Arnold and Mabel Beckman Center for CryoEM for cryo-EM support and the University of Utah Center for High Performance Computing for computational support. We thank Dr. Yorgo Modis for kindly sharing the assembled MDA5 trimer models of different nucleotide-bound states, which facilitated structural comparisons with our structure of the AVC. This work was supported by funding to B.L.B. (R35GM141262) and P.S.S. (R35GM133772) from the National Institute of General Medical Sciences of the NIH. P.J.N. is a Merck Fellow of the Damon Runyon Cancer Research Foundation (DRG-2564-25). B.L.B. is a Jon M. Huntsman Presidential Endowed Chair.

## Methods

### Protein expression and purification of DCR-1•DRH-1•RDE-4

Protein expression and purification were conducted as previously described^20^. Briefly, DCR-1, DRH-1, and RDE-4 were simultaneously co-expressed in *Spodoptera frugiperda* (Sf9) cells using the modified Bac-to-Bac baculovirus expression system. Harvested cells were resuspended in 5.5 times the pellet volume in buffer (25mM HEPES [pH 7.5]; 10mM KOAc; 2mM Mg(OAc)2; 100mM KCl; 1mM TCEP; 5% glycerol) and supplemented with 1% Triton X-100, 20 nM Avidin, 250 mg/ml DNase I Grade II, cOmplete EDTA-free protease inhibitor (1 tablet per 25 ml buffer), 0.7% (vol/vol) Protease Inhibitor Cocktail, and 1 mM phenylmethanesulfonyl fluoride (PMSF) prior to cell lysis. DCR-1•DRH-1•RDE-4 was purified in two liquid chromatography steps: StrepTrap HP affinity column followed by size-exclusion with a Superose 6 Increase 10/300 column.

### dsRNA Preparation

106 nucleotide (nt) RNAs were prepared using in vitro transcription as previously described ^29,47^. Equimolar amounts of ssRNAs were annealed in annealing buffer (50 mM TRIS pH 8.0, 20 mM KCl) by placing the reaction on a heat block (95°C) and slow cooling ≥2 hr.

### dsRNA Sequences

106 nt sense RNA: 5′- GGCAAUGAAAGACGGUGAGCUGGUGAUAUGGGAUAGUGUUCACCCUUGUUACACCGUUUUCCAUGAGCAAACUGAAACGUUUUCAUCGCUCUGGAGUGAAUACCAA-3′

106 nt antisense 3′ovr RNA: 5′- GGUAUUCACUCCAGAGCGAUGAAAACGUUUCAGUUUGCUCAUGGAAAACGGUGUAACAAGGGUGAACACUAUCCCAUAUCACCAGCUCACCGUCUUUCAUUGCCAA-3′

### Cryo-EM specimen preparation

1.2 µM DCR-1•DRH-1•RDE-4 with 1.2 µM 3’ovr dsRNA and with or without 5mM ATP was incubated at 20°C for 30 – 40 minutes in cleavage buffer (25mM HEPES [pH 7.5], 50mM KCl, 10mM Mg(OAc)_2_, 1mM TCEP). 3.5 μl of the reaction was applied to UltrAuFoil R2/2 Au300 mesh grids (Quantifoil) that were glow discharged for 25 seconds on each side at 25 mA using a Pelco easiGlow unit (Ted Pella, Inc.). The grids were blotted with filter paper (595 Filter Paper, Ted Pella, Inc.) for 1.5 seconds using an Mk. IV Vitrobot (Thermo Fisher Scientific) and then plunge frozen into liquid ethane.

### Cryo-EM and data processing of DCR-1•DRH-1•RDE-4 with 3’ovr dsRNA and 5mM ATP

Cryo-EM movies were recorded at a nominal magnification of 81,000x, corresponding to a pixel size of 1.058Å with a total dose of 50 electrons/Å^2^ and 50 frames per movie. Movies were patch motion corrected and CTF estimated in CryoSPARC v3.3.233^57^, and a total of 21,599 micrographs displaying CTF resolution fits of 8 Å or better were used for downstream analysis. Particles were selected using blob-based picking (150-250 Å diameter). After several rounds of 2D classification, a total of 2,617,679 particles were used for ab initio reconstruction (3 classes). After removing junk classes, a total of 1,639,881 particles were then used for heterogeneous refinement (4 classes). This led to one well-resolved class comprising 627,112 particles. The particles were subject to non-uniform refinement, which produced a 3.0 Å map. Data processing workflow is available in Figure S1.

### Cryo-EM and data processing of DCR-1•DRH-1•RDE-4 with 3’ovr dsRNA without ATP

Cryo-EM movies were recorded at a nominal magnification of 81,000x, corresponding to a pixel size of 1.058Å with a total dose of 50 electrons/Å^2^ and 50 frames per movie. Movies were patch motion corrected and CTF estimated in CryoSPARC v3.3.233^57^, and a total of 21,856 micrographs displaying CTF resolution fits of 6 Å or better were used for downstream analysis. Particles were selected using blob-based picking (150-250 Å diameter). After several rounds of 2D classification, a total of 171,294 particles were used for template-based picking, which found 7,085,361 particles. After several rounds of 2D classification, 1,152,125 particles were used for ab initio reconstructions (3 classes). The 1,152,125 particles were subjected to another round of 2D classification, where 486,581 particles selected. These particles and one of the ab initio classes were used for non-uniform refinement, which produced a 3.2 Å map. Topaz was then used to find additional particles that were combined with the 486,581 particles. These particles were subjected to the remove duplicates tool and resulted in 574,362 particles being used for non-uniform refinement that produced a 3.2 Å map that showed an improved density quality. Data processing workflow is available in Figure S5.

### Model Building, Refinement, and Validation

The model for DCR-1::DRH-1::RDE4::dsRNA with 5mM ATP was built manually in the 3.0 Å density map using Coot v0.8.9.1^58^. The AlphaFold2^59,60^ models of DCR-1 (AF-P34529-F1-v4), DRH-1 (AF-G5EDI8-F1-v4), and RDE-4 (AF-G5EBF5-F1-v4) were used as a starting point. We used de novo model building of the dsRNA, Mg^2+^, Zn, and ADP ligands. The dsRNA was built using A-form restraints in COOT. The model and substrates were subjected to real-space refinement using Phenix v1.20.1-4487. Default settings were used, except the weight was set to 0.01 and the nonbonded weight was set to 1500.

The model for DCR-1::DRH-1::RDE4::dsRNA without ATP was built manually in the 3.2 Å density map using Coot v0.8.9.1^58^. We used our plus ATP model as a starting point where we fit each protein into the new density individually. We then used the AlphaFold2 model of RDE-4 (AF-G5EBF5-F1-v4) to fit dsRBM1 and dsRBM2. We deleted the dsRNA from the starting model and rebuilt it from scratch using A-form dsRNA restraints in COOT. Although we saw additional density for dsRNA, we stopped building the dsRNA when the local resolution became too low to confidently build more nucleotides into the density. We note that our minus ATP density likely represents an average between no cleavage and one cleavage event, but favors a density without the dsRNA having been cleaved since when analyzed at the same thresholds of our plus ATP structure we see additional density for dsRNA (see Figure S6B). We also deleted the ADP ligands from our starting model as we did not see any density to support its presence, which confirmed we did not have any ATP contamination in our minus ATP dataset. The model and substrates were subjected to real-space refinement using Phenix v1.20.1-4487 and the same settings were used as in our structure that contained ATP.

