## Supplementary Information for "*C. elegans* Dicer stacks with the RIG-I-like receptor DRH-1 to cleave dsRNA"

| Supplementary Table: Cryo-EM data collection, refinement, and validation statistics | | |
|---|---|---|
| **Data collection and processing** | | |
| | Complex plus ATP | Complex minus ATP |
| EM Databank Accession ID | EMD-77587 | EMD-77700 |
| Microscope | Titan Krios G3 | Titan Krios G3 |
| Voltage (kV) | 300 | 300 |
| Detector | Gatan K3 | Gatan K3 |
| Data collection software | SerialEM | SerialEM |
| Nominal magnification | 81,000x | 81,000x |
| Total number of frames | 50 | 50 |
| Total electron exposure ($e^-$/Å) | 50 | 50 |
| Defocus range (µm) | -1.0 - -2.0 µm | -1.0 - -2.0 µm |
| Pixel size (Å) | 1.058 Å | 1.058 Å |
| Number of micrographs | 22,019 | 22,860 |
| Final particles | 627,112 | 574,362 |
| Resolution (GSFSC 0.143) | 3.0 Å | 3.2 Å |
| **Refinement and validation statistics for 2.9 Å reconstruction of DRH-1** | | |
| **Structure** | | |
| Protein Data Bank Accession ID | 36HV | 36NK |
| **Symmetry imposed** | | |
| 3D classification | C1 | C1 |
| 3D refinement | C1 | C1 |
| Final particles | 627,112 | 574,362 |
| **Map resolution (Å)** | | |
| FSC 0.143 (unmasked) | 3.5 Å | 3.5 Å |
| FSC 0.143 (masked, corrected) | 3.0 Å | 3.2 Å |
| **Model Refinement** | | |
| Initial model used (AlphaFold database) | DCR-1: AF-P34529-F1-v4<br>DRH-1: AF-G5EDI8-F1-v4<br>RDE-4: AF-G5EBF5-F1-v4 | |
| Map correlation coefficient | 0.85 | 0.87 |
| **Model composition** | | |
| Non-hydrogen atoms | 30147 | 31457 |
| Protein residues | 3298 | 3489 |
| Nucleotides | 166 | 160 |
| Ligands (ADP, $Mg^{2+}$, $Zn^{2+}$) | 12 | 6 |
| **R.m.s. deviations** | | |
| Bond lengths (Å) | 0.004 | 0.003 |
| Bond angles (°) | 0.856 | 0.661 |
| **Validation** | | |
| MolProbity score | 1.52 | 1.34 |
| Clashscore | 4.87 | 3.45 |
| Poor rotamers (%) | 0.07 | 0.03 |
| **Ramachandran plot** | | |
| Favored (%) | 96.12 | 96.73 |
| Allowed (%) | 3.88 | 3.27 |
| Disallowed (%) | 0.00 | 0.00 |
| C-beta deviations (0.25 Å) | 0.00 | 0.00 |
| CaBLAM outliers (%) | 2.16 | 1.95 |

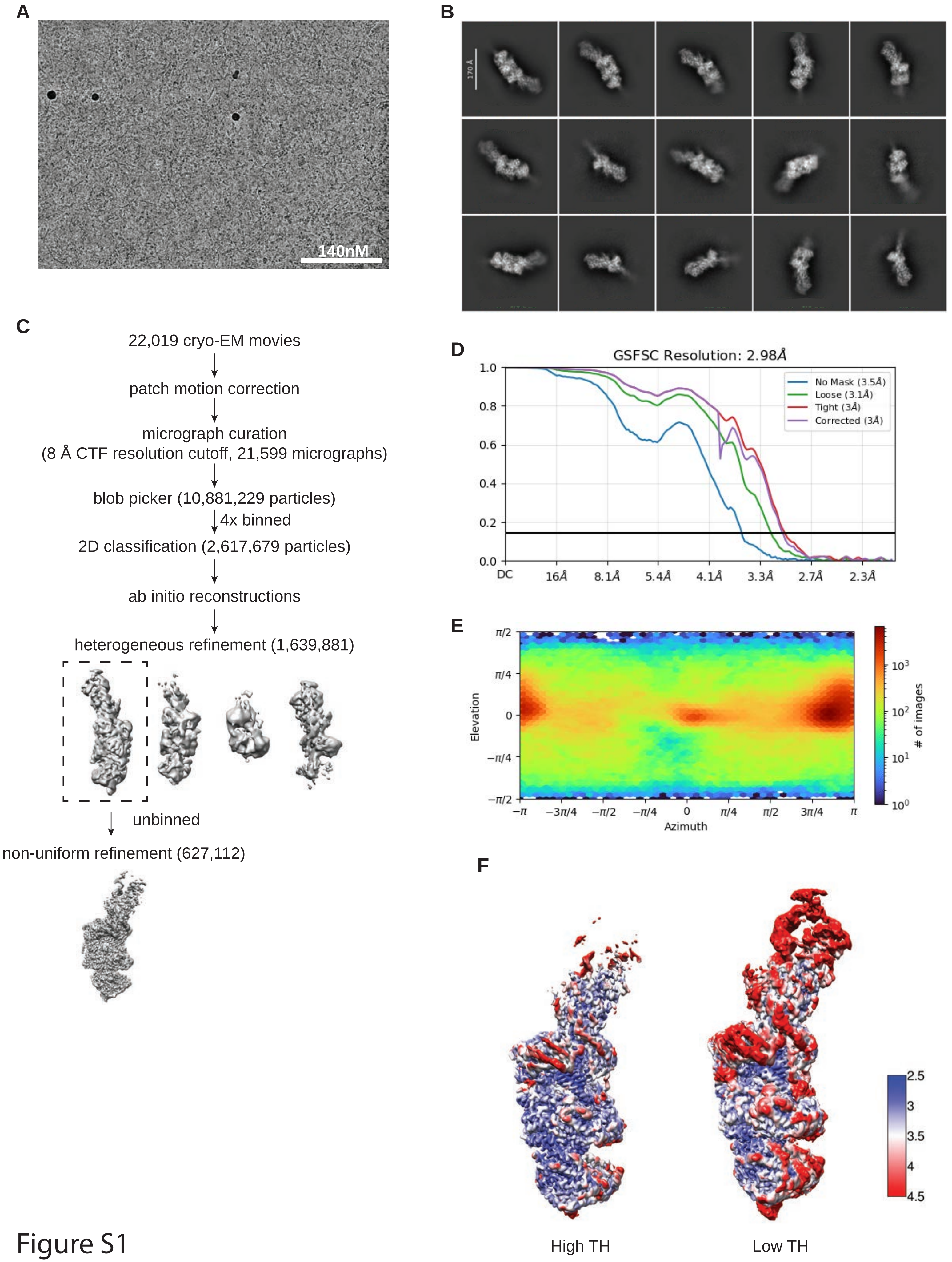
A
140nM
B
170 Å
C
22,019 cryo-EM movies
patch motion correction
micrograph curation
(8 Å CTF resolution cutoff, 21,599 micrographs)
blob picker (10,881,229 particles)
4x binned
2D classification (2,617,679 particles)
ab initio reconstructions
heterogeneous refinement (1,639,881)
unbinned
non-uniform refinement (627,112)
D
GSFSC Resolution: 2.98Å
No Mask (3.5Å)
Loose (3.1Å)
Tight (3Å)
Corrected (3Å)
DC
16Å
8.1Å
5.4Å
4.1Å
3.3Å
2.7Å
2.3Å
E
Elevation
Azimuth
### of images
F
2.5
3
3.5
4
4.5
High TH
Low TH

Figure S1

**Figure S1. Validation of the AVC structure with 3′ovr 106-dsRNA and ATP**

A. Representative cryo-EM micrograph.

B. Representative 2D class averages.

C. Processing tree of cryo-EM data in cryoSPARC.

D. Gold-standard FSC plot.

E. Orientation distribution of particles.

F. Local resolution heat map at high or low TH as indicated.

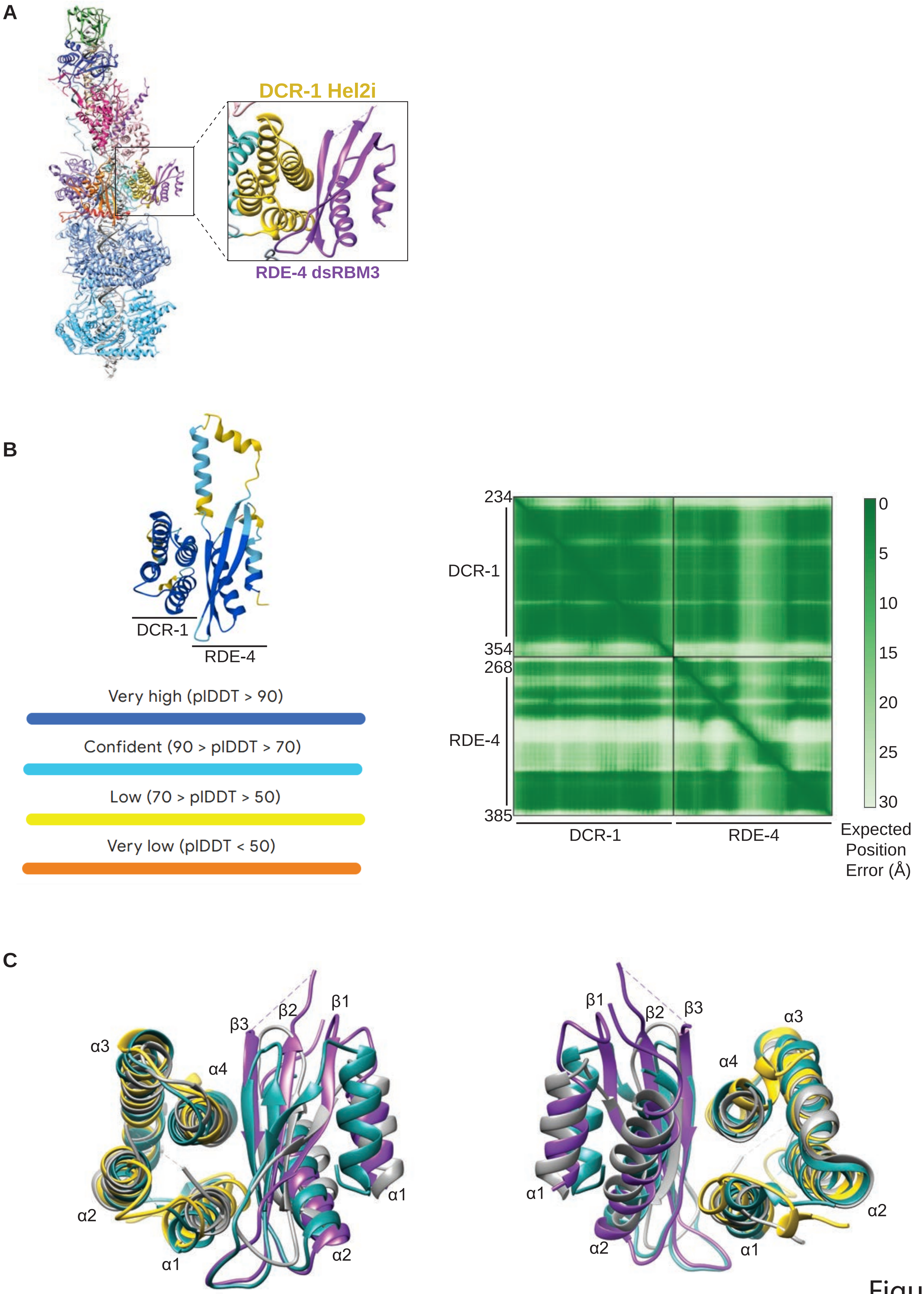
A
DCR-1 Hel2i
RDE-4 dsRBM3
B
DCR-1
RDE-4
Very high (plDDT > 90)
Confident (90 > plDDT > 70)
Low (70 > plDDT > 50)
Very low (plDDT < 50)
234
354
268
385
DCR-1
RDE-4
DCR-1
RDE-4
0
5
10
15
20
25
30
Expected Position Error (Å)
C
α3
α4
α2
α1
β3
β2
β1
α1
α2
β1
β2
β3
α3
α4
α1
α2
α2
α1

Figure S2

**Figure S2. AlphaFold prediction of interaction between DCR-1 Hel2i and RDE-4 third dsRBM.**

A. Magnified view of the interaction between DCR-1 Hel2i (yellow) and RDE-4 dsRBM3 (purple).

B. AlphaFold prediction of the *C. elegans* Hel2i and dsRBM3 interaction. Top: model of the interaction with confidence of tertiary structure fold as color-coded. Bottom: predicted aligned error (PAE) plot.

C. Two views of the structural alignment of the Hel2i and dsRBM3 interaction seen in *C. elegans* (colored as in A), human DCR-1•TRBP (5ZAM, dark cyan), and *Drosophila melanogaster* Dcr2•R2D2 (7V6C, gray).

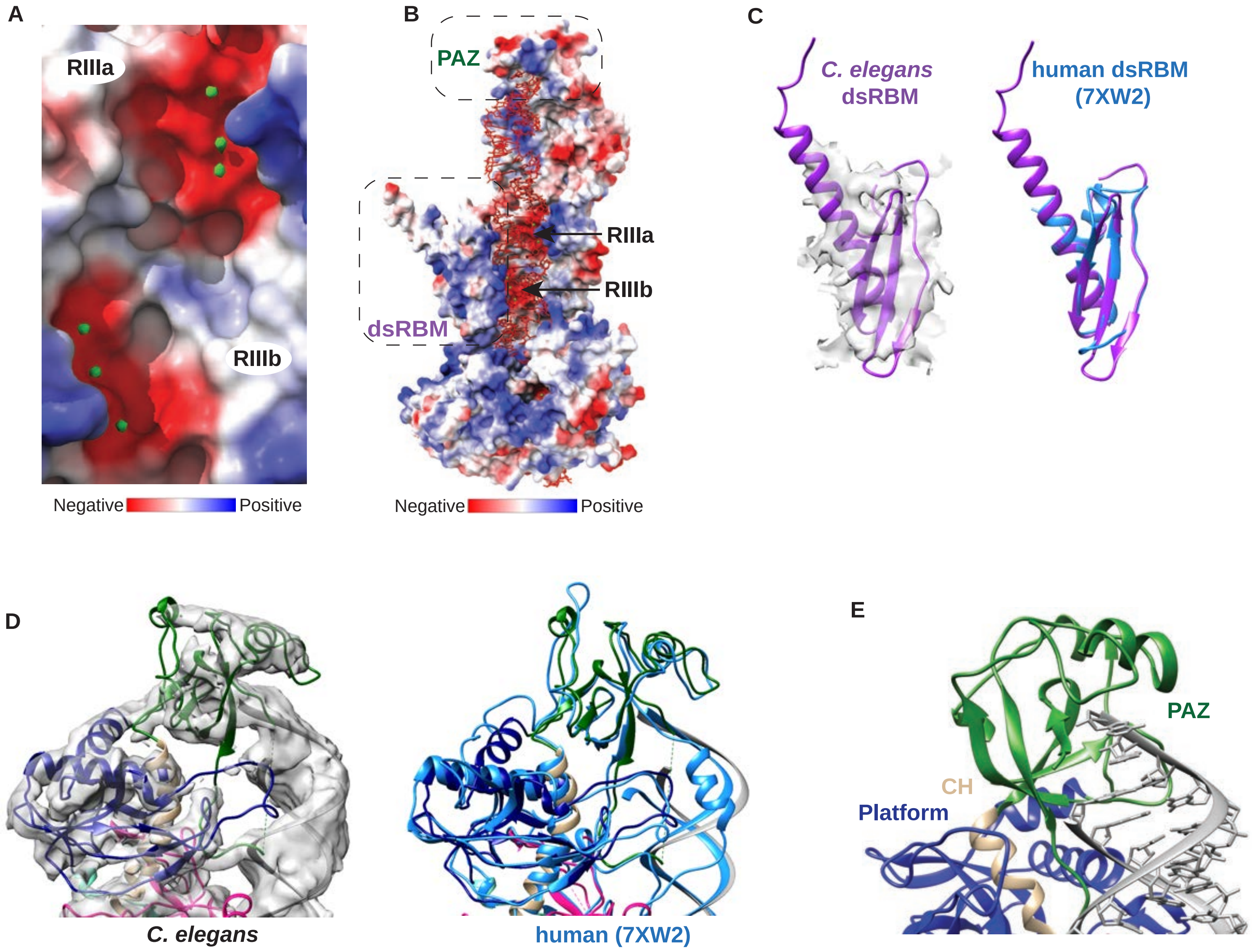
A
RIIIa
RIIIb
Negative
Positive
B
PAZ
RIIIa
RIIIb
dsRBM
Negative
Positive
C
C. elegans
dsRBM
human dsRBM
(7XW2)
D
C. elegans
human (7XW2)
E
PAZ
CH
Platform

Figure S3

**Figure S3. Dicer supplement**

A. Electrostatic potential surface model of DCR-1's RNase III domains. RIIIa (top) and RIIIb (bottom). Green spheres indicate $Mg^{+2}$ ions positioned in the RIII negatively charged electrostatic pockets.

B. Electrostatic potential surface model of Dicer with the following domains indicated: RIIIa, RIIIb, PAZ, and dsRBM.

C. Left: *C. elegans* DCR-1 dsRBM (purple) overlayed with the density map. Right: Structural alignment between *C. elegans* (purple) and human (7XW2, blue) Dicer dsRBMs.

D. Left: *C. elegans* PAZ (green), Platform (dark blue), and dsRNA (gray) overlayed with the density map. Right: Structural alignment between *C. elegans* and human (7XW2, light blue).

E. Magnified view of the 2-nucleotide 3'overhang extending into the PAZ domain.

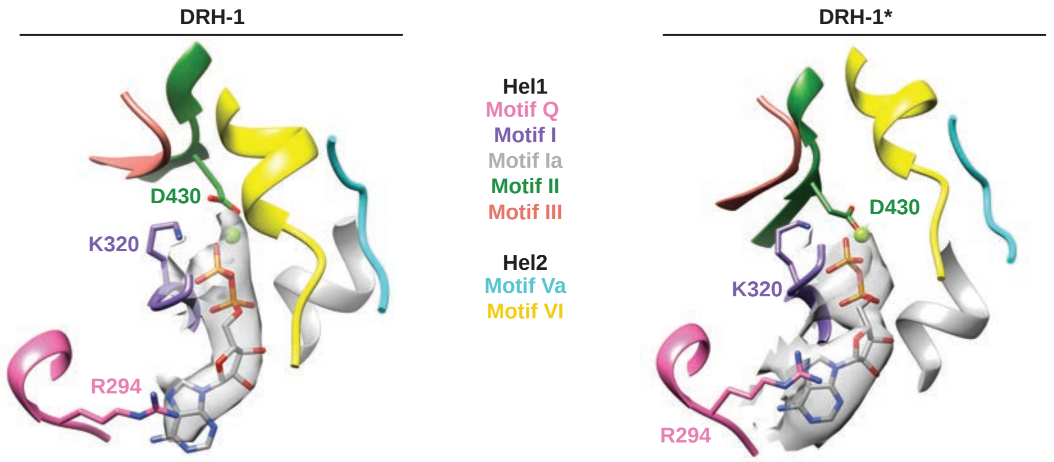
DRH-1
DRH-1*
Hel1
Motif Q
Motif I
Motif Ia
Motif II
Motif III
Hel2
Motif Va
Motif VI
D430
K320
R294
D430
K320
R294

Figure S4

**Figure S4. ADP observed in DRH-1 and DRH-1* ATP-binding pockets.**

ADP and $Mg^{2+}$ (light green sphere) are overlayed with the density map and surrounding helicase motifs in DRH-1 and DRH-1*.

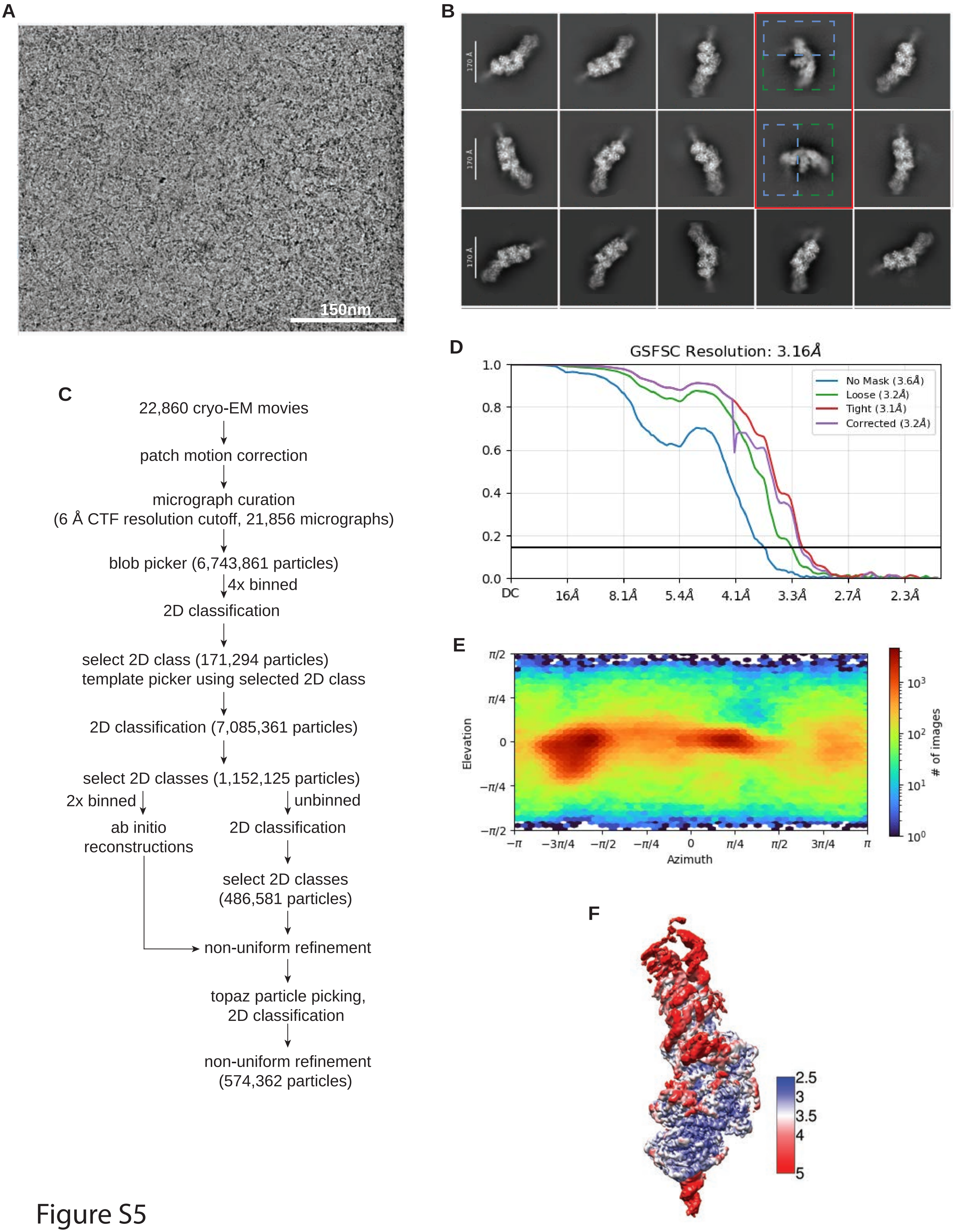
A
150nm
B
170 Å
170 Å
170 Å
C
22,860 cryo-EM movies
patch motion correction
micrograph curation
(6 Å CTF resolution cutoff, 21,856 micrographs)
blob picker (6,743,861 particles)
4x binned
2D classification
select 2D class (171,294 particles)
template picker using selected 2D class
2D classification (7,085,361 particles)
select 2D classes (1,152,125 particles)
2x binned
unbinned
ab initio
reconstructions
2D classification
select 2D classes
(486,581 particles)
non-uniform refinement
topaz particle picking,
2D classification
non-uniform refinement
(574,362 particles)
D
GSFSC Resolution: 3.16Å
No Mask (3.6Å)
Loose (3.2Å)
Tight (3.1Å)
Corrected (3.2Å)
1.0
0.8
0.6
0.4
0.2
0.0
DC
16Å
8.1Å
5.4Å
4.1Å
3.3Å
2.7Å
2.3Å
E
π/2
π/4
0
−π/4
−π/2
Elevation
−π
−3π/4
−π/2
−π/4
0
π/4
π/2
3π/4
π
Azimuth
10³
10²
10¹
10⁰
### of images
F
2.5
3
3.5
4
5

Figure S5

**Figure S5. Validation of the AVC structure with 3′ovr 106-dsRNA.**

A. Representative cryo-EM micrograph.

B. Representative 2D class averages. Red box indicates 2D classes that represent the complex in an initial binding state where DCR-1 is denoted in a green dashed box and DRH-1 is indicated with a blue dashed box.

C. Processing tree of cryo-EM data in cryoSPARC.

D. Gold-standard FSC plot.

E. Orientation distribution of particles.

F. Local resolution heat map.

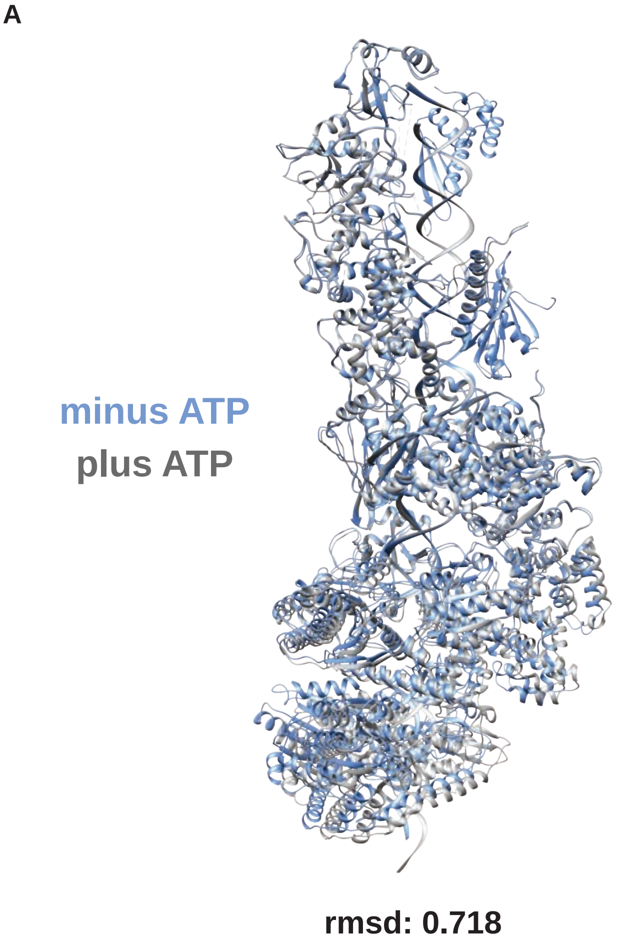
A
minus ATP
plus ATP
rmsd: 0.718

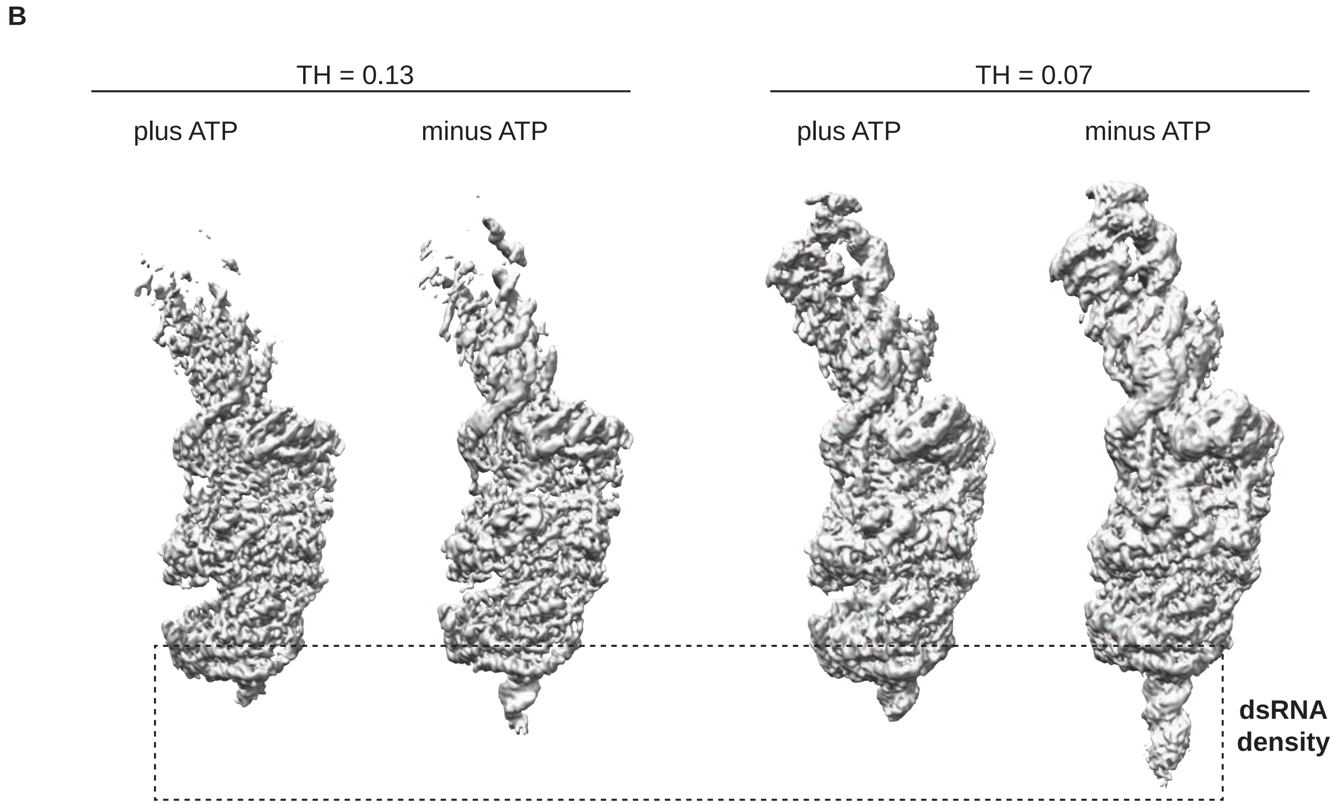
B
TH = 0.13
TH = 0.07
plus ATP
minus ATP
plus ATP
minus ATP
dsRNA
density

Figure S6

**Figure S6.**

A. Structural alignment of the plus (gray) and minus (blue) ATP models.

B. Densities of minus and plus ATP with different thresholds displayed as indicated. The differences in dsRNA densities are outlined in a dashed box.

A

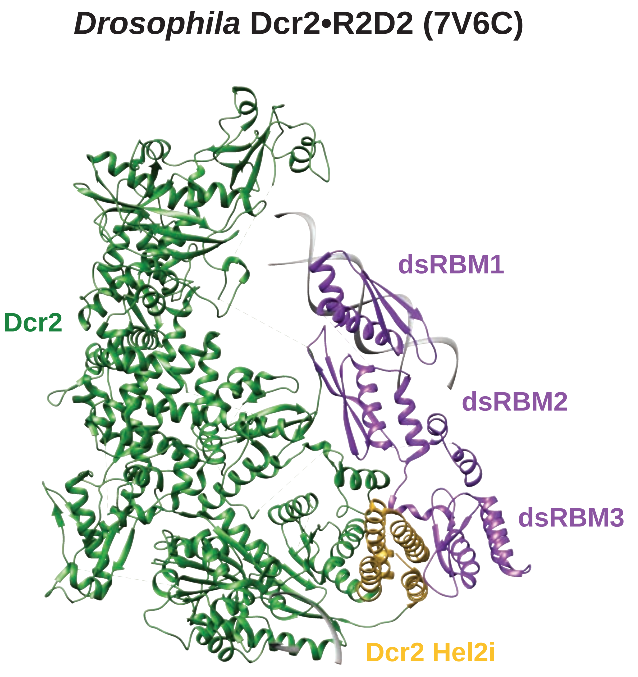

*Drosophila* Dcr2•R2D2 (7V6C)
Dcr2
dsRBM1
dsRBM2
dsRBM3
Dcr2 Hel2i


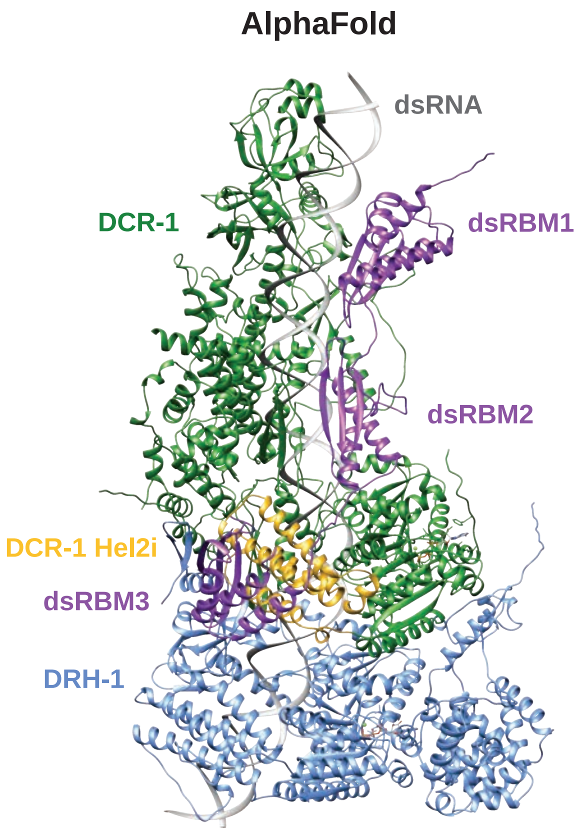

AlphaFold
dsRNA
DCR-1
dsRBM1
dsRBM2
DCR-1 Hel2i
dsRBM3
DRH-1


Figure S7

**Figure S7.**

A. Left: Model of Drosophila Dcr2•R2D2 (7V6C). Right: AlphaFold prediction of the AVC with blunt 106-dsRNA, two ATP molecules, and two $Mg^{2+}$ ions. Proteins and domains are colored as indicated.

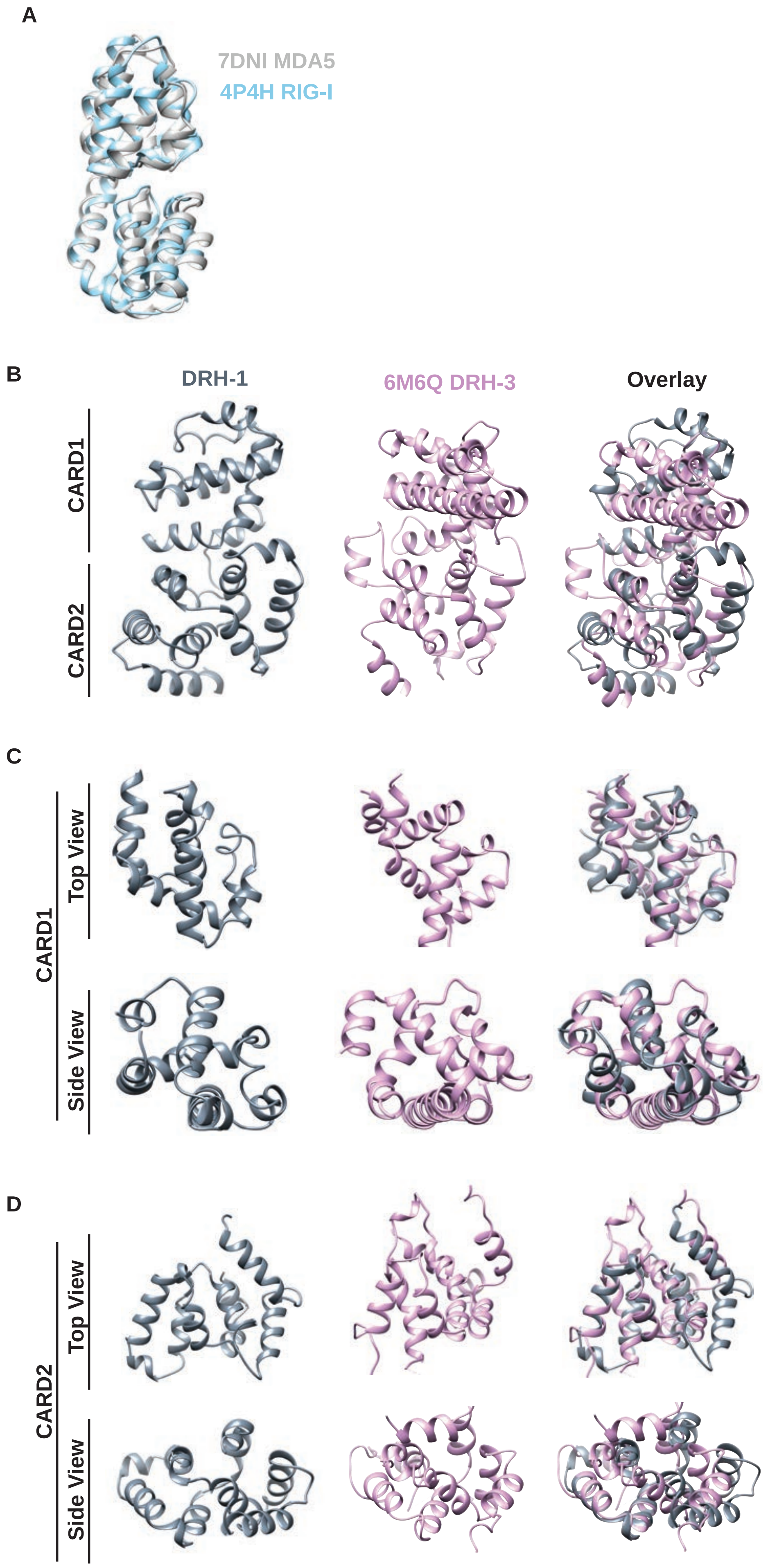
A
7DNI MDA5
4P4H RIG-I
B
DRH-1
6M6Q DRH-3
Overlay
CARD1
CARD2
C
CARD1
Top View
Side View
D
CARD2
Top View
Side View

Figure S8

**Figure S8.**

A. Structural alignment between MDA5 (7DNI, gray) and RIG-I (4P4H, blue) CARDs.

B. Structure of the DRH-1 NTD (left), DRH-3 NTD (6M6Q, middle), and overlay between both (right). Structures are oriented to show the NTD with CARD1 above and CARD2 below.

C. Top and side views of CARD1 from DRH-1 (left) and DRH-3 (6M6Q, middle), and an overlay between both.

D. Top and side views of CARD2 from DRH-1 (left) and DRH-3 (6M6Q, middle), and an overlay between both.

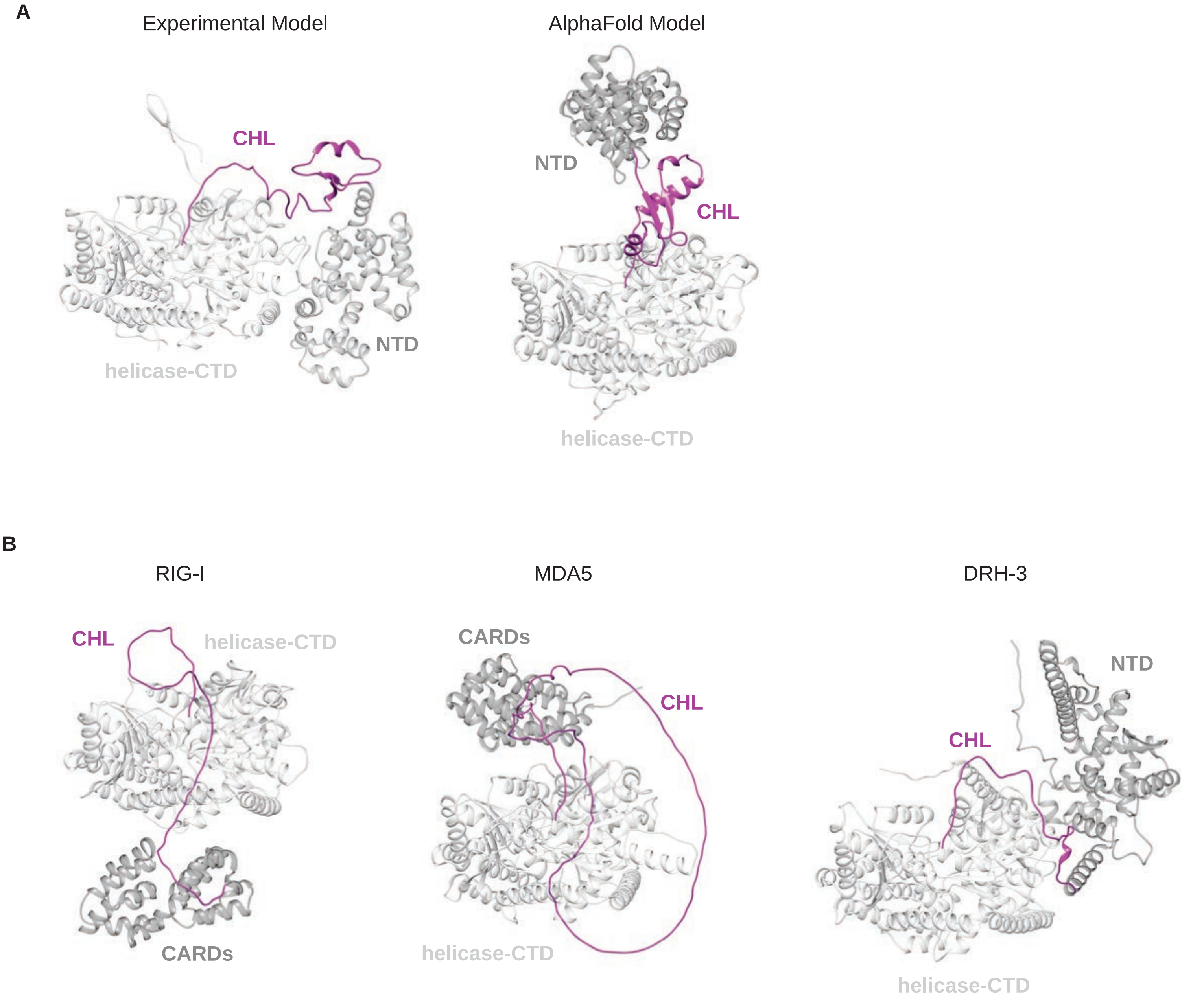
A
Experimental Model
AlphaFold Model
CHL
NTD
helicase-CTD
NTD
CHL
helicase-CTD
B
RIG-I
MDA5
DRH-3
CHL
helicase-CTD
CARDs
CARDs
CHL
helicase-CTD
NTD
CHL
helicase-CTD

Figure S9

**Figure S9. RLRs models.**

A. Experimental and AlphaFold models of DRH-1 highlighting the structured CHL colored in magenta. Both models are shown with the helicase in the same orientation.

B. AlphaFold models of RIG, MDA5, and DRH-3 highlighting the unstructured helicase linker colored in magenta. All models are shown with the helicase in the same orientation.

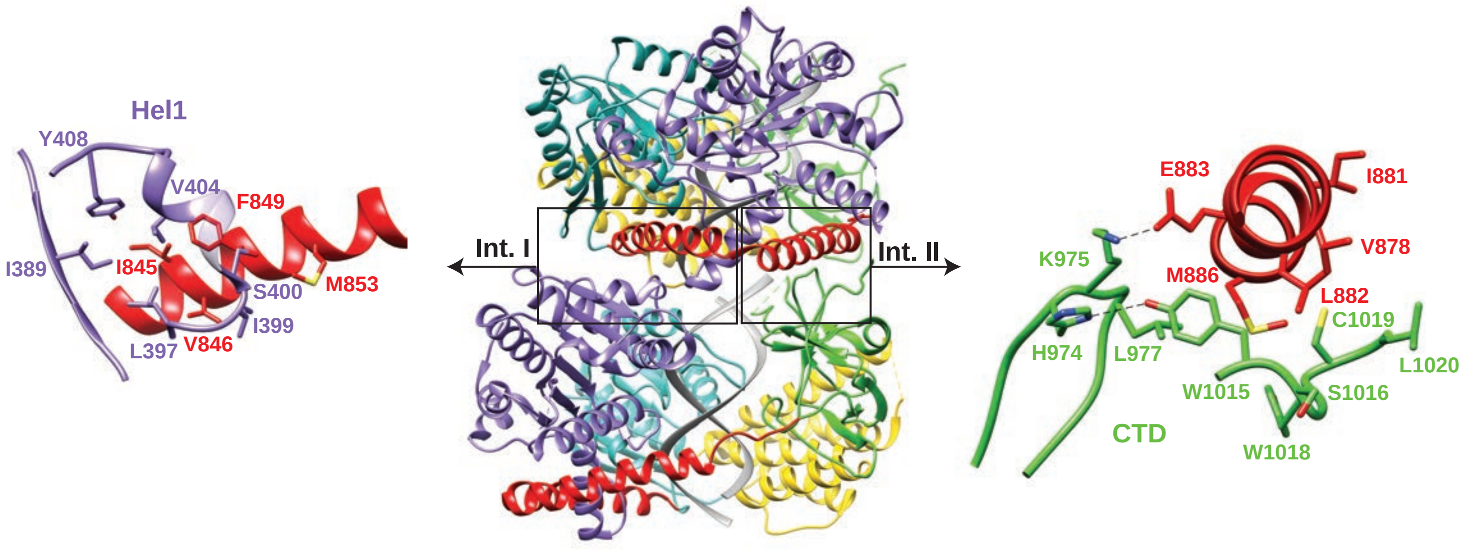

Hel1
Y408
V404
F849
I389
I845
S400
M853
I399
L397
V846
Int. I
Int. II
E883
I881
K975
V878
M886
L882
C1019
H974
L977
L1020
W1015
S1016
CTD
W1018


Figure S10

**Figure S10. MDA5 stacking interactions are mediated by hydrophobic and electrostatic interactions.**

MDA5 monomers (PDB 6G19) were fit into the EMD-4338 density map to build the stacking interaction as discovered and described in *Motis et al*[39] . The domains are colored as shown in Figure 1A.

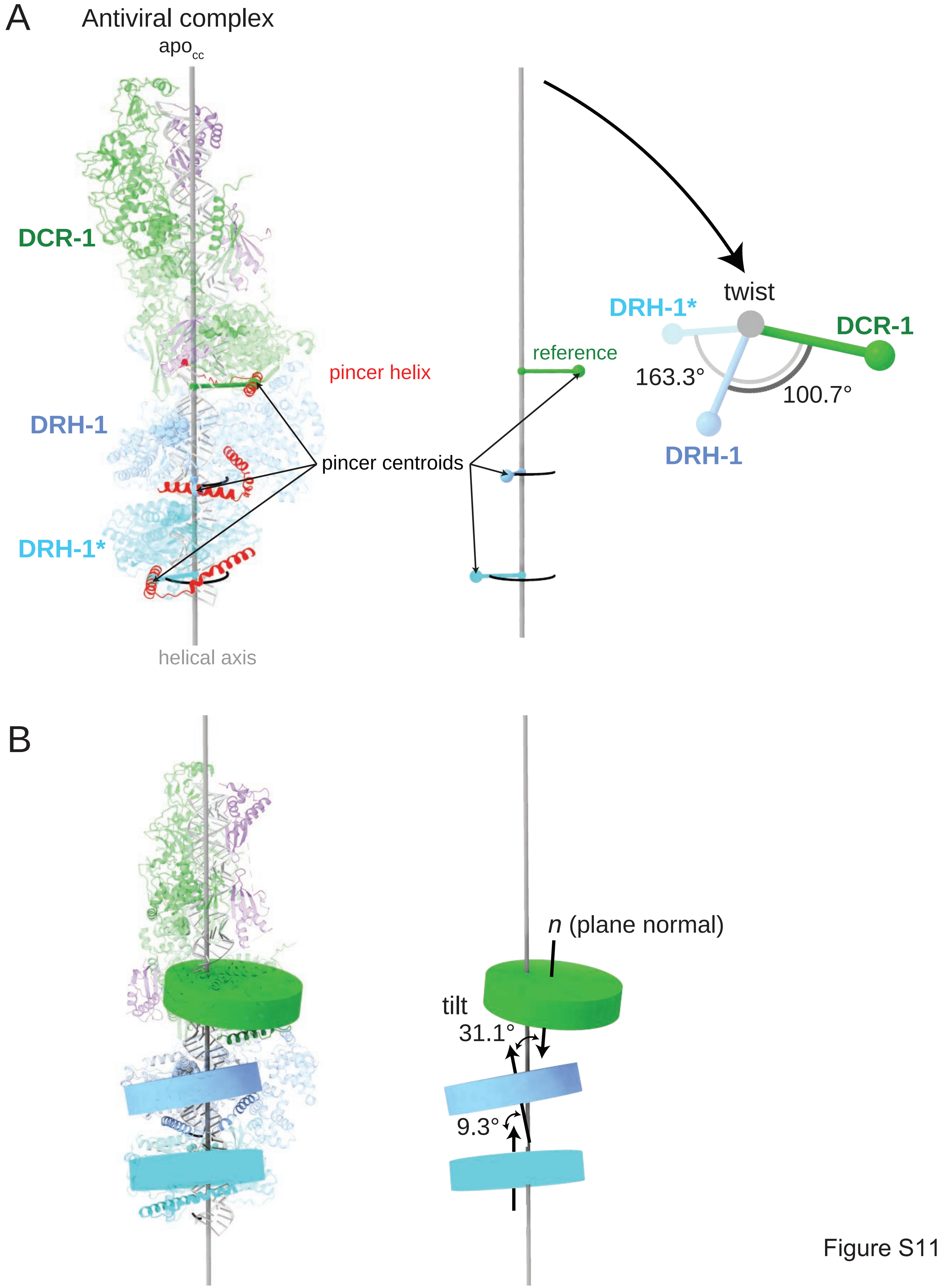
A
Antiviral complex
apo$_{cc}$
DCR-1
DRH-1
DRH-1*
pincer helix
pincer centroids
helical axis
reference
twist
DRH-1*
DCR-1
163.3°
100.7°
DRH-1
B
n (plane normal)
tilt
31.1°
9.3°

Figure S11

**Figure S11. Method for measurement of twist and tilt between adjacent helicases in the antiviral complex and MDA5 trimer.**

A. Helical twist between adjacent helicase domains was measured relative to the central dsRNA axis by projecting a shared pincer-domain (red) centroid onto the RNA axis and calculating the angular displacement between neighboring monomers.

B. Inter-subunit tilt was quantified by fitting a best-fit plane to the Cα atoms of each helicase core and calculating the angle between the corresponding plane normals.

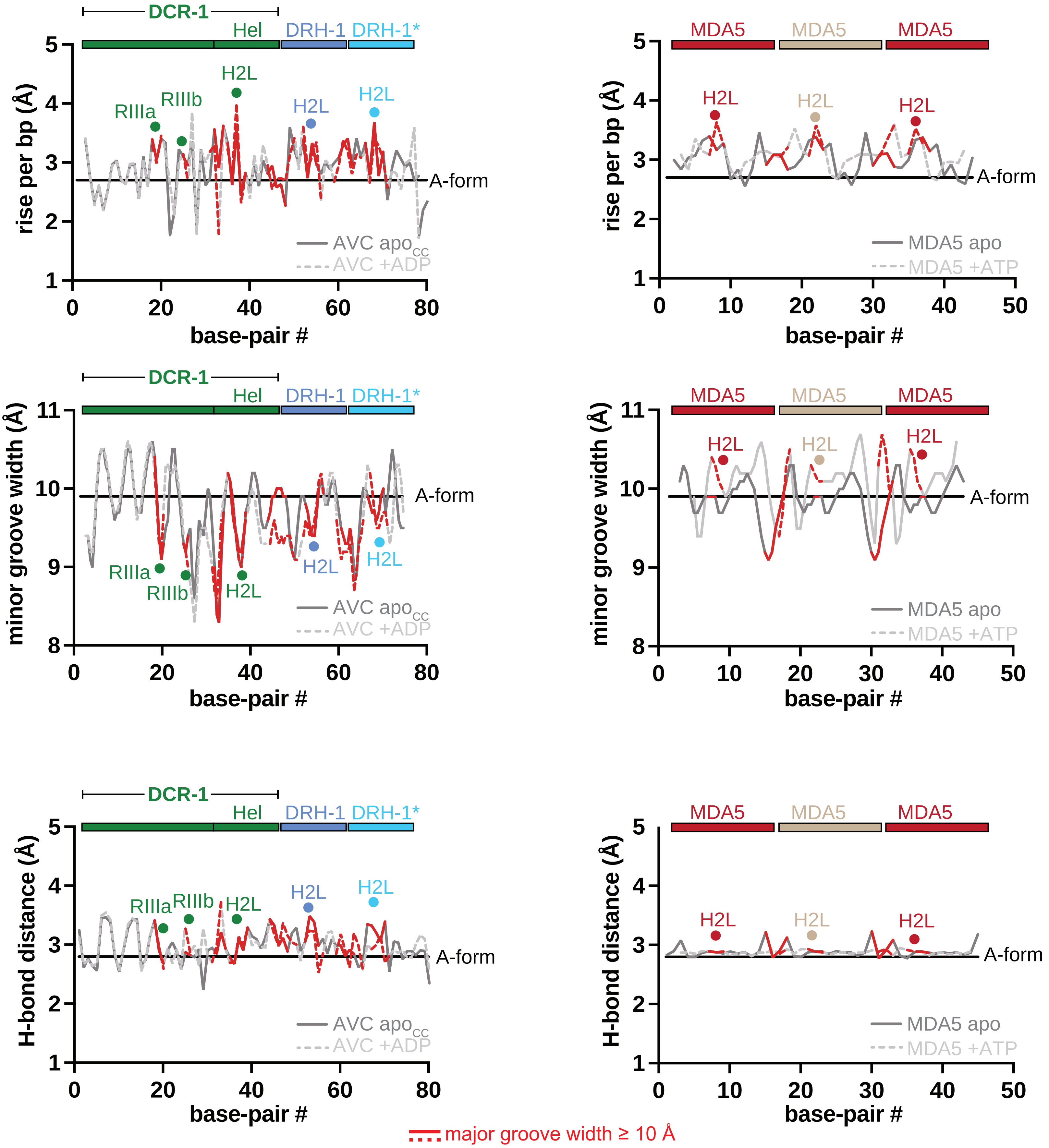

DCR-1
Hel
DRH-1
DRH-1*
MDA5
H2L
RIIIa
RIIIb
A-form
rise per bp (Å)
minor groove width (Å)
H-bond distance (Å)
base-pair #
AVC apo$_{CC}$
AVC +ADP
MDA5 apo
MDA5 +ATP
major groove width ≥ 10 Å

**Figure S12. Distortion of dsRNA away from A-form by AVC and MDA5 binding.**

Graphs of rise per bp, minor groove widths, and basepair H-bond distances (N1-N3) for A-form dsRNA and the dsRNA associated with the $apo_{CC}$ and ADP-bound AVC and ATP-bound MDA5 trimer models. The lines are colored red according to the regions of major groove widening ≥10Å as shown in Figure 6A,6B. The locations of the RNase IIIa/b catalytic residues, Hel2 loops, and interfaces between helicase domains are indicated as dots above the lines. The location of the domains of the antiviral complex and MDA5 trimer relative to the dsRNA is indicated above the plots.
